# Maintenance of quantitative genetic variance in complex, multi-trait phenotypes: The contribution of rare, large effect variants in two Drosophila species

**DOI:** 10.1101/2022.04.21.488876

**Authors:** Emma Hine, Daniel E. Runcie, Scott L. Allen, Yiguan Wang, Stephen F. Chenoweth, Mark W. Blows, Katrina McGuigan

## Abstract

The interaction of evolutionary processes to determine quantitative genetic variation has implications for contemporary and future phenotypic evolution, as well as for our ability to detect causal genetic variants. While theoretical studies have provided robust predictions to discriminate among competing models, empirical assessment of these has been limited. In particular, theory highlights the importance of pleiotropy in resolving observations of selection and mutation, but empirical investigations have typically been limited to few traits. Here, we applied high dimensional Bayesian Sparse Factor Genetic modelling to 3,385 gene expression traits from *Drosophila melanogaster* and from *D. serrata* to explore how genetic variance is distributed across high-dimensional phenotypic space. Surprisingly, most of the heritable trait covariation was due to few lines (genotypes) with extreme (>3 IQR from the median) values. This observation, in the two independently sampled species, suggests that the House of Cards (HoC) model might apply not only to individual expression traits, but also to emergent co-expression phenotypes. Intriguingly, while genotypes extreme for a multivariate factor also tended to have a higher proportion of individual traits that were extreme, we also observed genotypes that were outliers for multivariate factors but not for any individual traits. We observed other consistent differences between heritable multivariate factors with outlier lines versus those factors that conformed to a Gaussian distribution of genetic effects, including differences in gene functions. We use these observations to identify further data required to advance our understanding of the evolutionary dynamics and nature of standing genetic variation for quantitative traits.

## Introduction

The maintenance of quantitative genetic variance presents geneticists and evolutionary biologists with a formidable challenge. While models of the evolution of allele frequencies can be relatively simple, evolution of the genetic variance for phenotypic traits also depends on the effects of those alleles (Walsh and Lynch 2018). A substantial and complex body of theory has resulted in competing models, with no clear resolution of how quantitative variation evolves (Bürger 2000; Johnson and Barton 2005; Walsh and Lynch 2018).

Despite the central importance of the nature of quantitative genetic variance both for predicting long-term phenotypic evolution (Arnold *et al*. 2008), and for optimising approaches to identify causal loci (Eyre-Walker 2010; Simons *et al*. 2018), we have only limited empirical knowledge of the joint distribution of allele frequencies and their effects on traits of interest and on fitness (Johnson and Barton 2005; Walsh and Lynch 2018). Our purpose in this study is to revisit the basic observation of the distribution of genetic variance in complex quantitative phenotypes to help distinguish between the potential mechanisms underlying the maintenance of genetic variance. Our focus is on two key aspects of theoretical models: the relationship between allele frequency and effect size, and pleiotropy.

From a theoretical perspective, the resulting allele frequency spectrum is a key distinguishing feature between models in which selection actively maintains polymorphisms (balancing selection) and models in which selection eliminates variation (and mutation reintroduces it: mutation-selection balance, MSB). Balancing selection mechanisms are predicted to maintain relatively symmetrical allele frequencies at a locus, while under MSB models, genetic variation is determined by rare alleles, where the greater the fitness effect of a locus, the rarer the minor allele at that locus (Johnson and Barton 2005; Walsh and Lynch 2018).

Genomic studies of adaptation have provided data consistent with balancing selection models, such as fluctuation of allele frequencies with short-term environmental variation (e.g., Bergland *et al*. 2014), and the observation that common, not rare, alleles contribute to rapid adaptation (e.g., Kelly and Hughes 2019). On the other hand, large-scale genetic mapping studies in humans (Kemper *et al*. 2012; Zhao *et al*. 2016; Hernandez *et al*. 2019; Schoech *et al*. 2019) and other taxa (Josephs *et al*. 2015; Bloom *et al*. 2019) suggest a strong contribution to standing genetic variance of rarer alleles with relatively larger effects. These latter observations are consistent with MSB model predictions that selection acts to eliminate variation and is most effective at reducing the frequency of large effect alleles, causing a negative association between effect size and frequency.

Two different classes of MSB models make further contrasting predictions about allelic effect sizes based on the relative roles of mutation and selection in maintaining genetic variation. The house-of-cards (HoC) mutation model (Turelli 1984; Turelli 1985), where all mutations affect nearly-optimal alleles and are therefore deleterious, predicts selection is stronger than mutation, and that most standing genetic variance at each locus becomes a consequence of rare alleles of large effect (Walsh and Lynch 2018). In contrast, under the alternative Gaussian mutation model (Lande 1975) the normal (Gaussian) distribution of allelic effects at each locus results in mutation contributing more strongly than selection (Walsh and Lynch 2018), and genetic variance is due to many rare alleles of smaller effect. Humans GWAS of various phenotypes suggest that genetic variance is due to additive effects of many loci of small effect (reviewed in Simons *et al*. 2018), consistent with the Guassian model. In a rare example of an explicit test of predictions of HoC versus Guassian models, Hodgins-Davis et. al. (2015) found strong support for the HoC model for gene expression traits across three taxa. Other analyses of gene expression data are also consistent with HoC models, where the proportion of heritable variation explained by rare alleles indicates that their effect sizes must be much larger than the effects of more common alleles (Kremling *et al*. 2018; Hernandez *et al*. 2019). Mutation accumulation (MA) experiments have also provided evidence consistent with predictions of HoC models, reporting that most new mutations contribute little phenotypic variation, with few (rare) mutations having large phenotypic effects (Mackay *et al*. 1992; Davies *et al*. 1999; Heilbron *et al*. 2014; Mcguigan *et al*. 2014b).

Notably, empirical observations of selection, mutation and genetic variance are seemingly incompatible with any quantitative genetic theory on a trait-by-trait basis, leading to the incorporation of pleiotropy into theoretical models (Johnson and Barton 2005; Walsh and Blows 2009; Walsh and Lynch 2018). Although the empirical evidence for pleiotropy has been controversial (Paaby and Rockman 2013), advances in accessibility of genomic data, coupled with extensive phenotypic data, are revealing pleiotropic variants across diverse traits (Bulik-Sullivan *et al*. 2015; Chesmore *et al*. 2018; Geiler-Samerotte *et al*. 2020; Shikov *et al*. 2020). The empirical distribution of genetic variance in multiple traits presents a very different perspective on the maintenance of genetic variance than apparent when considering traits individually. While genetic variance in single traits appears essentially ubiquitous (Mousseau and Roff 1987; Houle 1992; Blows and Hoffmann 2005), most of the genetic variance in sets of multiple traits is typically restricted to a smaller subspace, defined by linear combinations of the measured traits (Kirkpatrick 2009; Walsh and Blows 2009; Blows and Mcguigan 2015). This uneven empirical distribution of genetic variance across phenotypic space implies that the number of genetically independent traits (*n*) is much lower than the number of traits measured (*p*). Indeed, based on genetic load and genetic variance in fitness arguments, *n* > 200 is predicted to be unlikely (Barton 1990; Johnson and Barton 2005).

High dimensional (*p* >> 10) genetic analyses to explicitly test Barton’s conjecture are rare, partly reflecting challenges of measuring many phenotypes on the same individual (genotype), and partly due to the statistical difficulties in accommodating large numbers of traits within a quantitative genetic framework. One solution to this dual challenge is the application of Bayesian sparse factor models to gene expression data (Runcie and Mukherjee 2013). This analytical approach has been used to show that, consistent with the expectation that *n* << *p*, much of the standing genetic (Runcie and Mukherjee 2013; Siren *et al*. 2017) and mutational (Hine *et al*. 2018) variance, can be explained by few linear combinations of the measured variables. Although these results support the conjecture that understanding pleiotropy is the key to understanding the maintenance of genetic variation (Johnson and Barton 2005), different theoretical models incorporating pleiotropy make very different predictions (Walsh and Lynch 2018), indicating that further empirical work will be important in helping to define a way forward.

Here, we aim to provide new insights into the nature of standing genetic variation by considering the relationship between variant frequency and effect size underpinning genetic covariances among traits. Theoretical models differ in the assumptions about the correlation of pleiotropic effect sizes among traits, specifically whether they are uncorrelated, or whether individuals that carry a pleiotropic allele that generates an extreme value for one trait will also be extreme for other traits (Turelli 1985; Barton 1990; Wingreen *et al*. 2003; Johnson and Barton 2005; Waxman and Peck 2006). While there is some evidence of stronger selection on alleles with highly pleiotropic effects (Denver *et al*. 2005; Mcguigan *et al*. 2014a), the distributions of allele frequency and of pleiotropic effects remains poorly characterised for any traits. Notably, theoretical models of MSB typically presume that mutations, while having negative effects on fitness, have unbiased effects on phenotypic traits (Johnson and Barton 2005), but the emergence of extreme multivariate trait values from the pleiotropic effects on each individual trait is unknown. Here, we ask whether the effects that genetic variants have on individual trait variation is predictive of the multivariate distribution of their effects across many traits, and whether the multivariate distribution of heritable phenotypes is more consistent with Gaussian or HoC predictions.

To investigate these questions, we use high dimensional Bayesian Sparse Factor Genetic (BSFG) modelling (Runcie and Mukherjee 2013) to interrogate the distribution of standing genetic variance in two unrelated datasets, one from *Drosophila serrata* (Allen *et al*. 2013; Mcguigan *et al*. 2014b) and one from *Drosophila melanogaster* (Ayroles *et al*. 2009; Runcie and Mukherjee 2013). In each case, we conduct the analysis on 3,385 gene expression traits, with 30 inbred lines capturing standing genetic variance in the natural population from which flies were sampled. There are three key parameters estimated in the BSFG model that form the basis of our investigations: the trait loadings of each (latent) phenotypic common factor (PCF), heritability of each PCF, and the latent trait values (PCF scores). The trait loadings of a given PCF allow us to determine how many expression traits are influenced by the putative latent factor (i.e., the extent of trait covariance), while the PCF heritability allows us to determine whether the latent factor is genetic or environmental. For both *D. serrata* and *D. melanogaster*, we confirm previous inferences of substantial genetic covariance of these gene expression traits (Ayroles *et al*. 2009; Runcie and Mukherjee 2013; Blows *et al*. 2015). The distribution of PCF scores across our fly lines then allowed us to infer the presence, in both species, of both genetic covariance due to common, small-effect variants, and genetic covariance due to rare, large-effect variants. Finally, we demonstrate that observed rare variants with effects on many traits are not caused by physically linked loci. This study provides evidence that the standing genetic covariance of expression traits is largely determined by rare alleles of major and pleiotropic effect.

## Methods

### Drosophila serrata dataset

The *D. serrata* inbred lines, and the characterisation of their gene expression, have been described in detail elsewhere (Allen *et al*. 2013; Mcguigan *et al*. 2014b). Briefly, iso-female lines were established from a natural, outbred population in St Lucia, Brisbane, Australia, and each line was subjected to 15 generations of full-sib mating (Allen *et al*. 2013). Gene expression was measured for two biological replicates of males from each of 30 iso-female lines, using a microarray approach. For each biological replicate, expression was measured using five different probes, each appearing twice on an individual array. For each probe, we calculated the mean log_10_ expression of the two replicates then calculated the median value across the five probes. The variance-standardized median expression was then analysed using the BSFG model. Most (75%) of the 11,604 assayed expression traits displayed significant genetic variance (5% false discovery rate), with an average broad-sense heritability of 0.34 (Mcguigan *et al*. 2014b). An earlier attempt at a high-dimensional analysis of these data constructed an 8,750-trait genetic covariance matrix from 175 50-trait matrices, where each 50-trait matrix was constructed by bivariate analysis of all pairs of traits; this analysis was statistically limited to consider only one multivariate axis of trait variation (Blows *et al*. 2015). We now use Runcie and Mukherjee’s (2013) BSFG modelling to fully characterise genetic covariance in this data. We restrict our analysis to a subset of 3,385 traits for which we have previously analysed mutational variance using the BSFG model (Hine *et al*. 2018).

### Drosophila melanogaster dataset

A set of 40 highly inbred lines were derived by 20 generations of full-sib mating from a natural *D. melanogaster* population in Raleigh, North Carolina, USA, as detailed in Ayroles *et al*. (2009). To facilitate comparability of the *D. serrata* and *D. melanogaster* datasets, we matched the dimensions of the *D. melanogaster* data to the *D. serrata* data, taking a random subset of 30 of the 40 inbred *D. melanogaster* lines, and a random subset of 3,385 of the 10,096 genetically variable gene expression traits. We considered only male data, analysing the two biological replicates per line, as for the *D. serrata* data. Previous analysis of these data indicated strong patterns of genetic covariance among expression traits, with 241 modules identified through modulated modularity clustering (Stone and Ayroles 2009), including two large sets of covarying expression traits (Ayroles *et al*. 2009). Runcie and Mukherjee (2013) previously subjected a much smaller subset (414) of these traits, implicated as being involved in competitive fitness, to BSFG modelling.

### Statistical Analysis

We employed the Bayesian Sparse Factor Genetic model (BSFG) described by Runcie and Mukherjee (2013) to partition the variance among and within lines. Briefly, the model is:

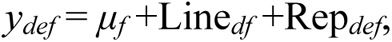

where *y_def_* is the measurement of gene *f* for the *e*^th^ replicate of line *d*, *μ_f_* is the global mean for gene *f*, *Line_df_* is the mean effect of line *d* on gene *f* and *Rep_def_* is the mean effect for replicate *e* on gene *f*. Line and replicate effects are considered random with distributions MVN*_F_*(0,**G**) and MVN*_F_*(0,**R**), respectively. Covariance matrices, **G** and **R**, are modelled as:

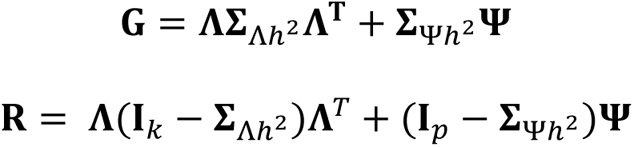

where *k* is the number of phenotypic common factors (PCFs) determined by the model, *p* is the number of gene expression traits (here, 3,385), **Λ** is a *p* × *k* matrix of factor loadings, **∑**_Λh^2^_ is a *k* × *k* diagonal matrix of PCF heritabilities, **Ψ** is a *p* × *p* diagonal matrix of trait-specific variances, **∑**_Ψh^2^_ is the *p* × *p* diagonal matrix of heritabilities for the trait-specific variances and **I** is the *p* × *p* identity matrix.

The model was implemented using the BSFG package in R (Runcie 2021) using the same prior distribution hyperparameter values relating to PCF sparsity and trait loading magnitude for both datasets. These hyperparameter values were the same as those used in the previous application of the model, implemented in MATLAB, on the 3,385 *D. serrata* gene expression traits in mutation accumulation lines (see Table S1 Hine *et al*. 2018). We specified the model to estimate trait loadings and heritability for 59 PCFs, the maximum possible non-zero eigenvalues in the phenotypic covariance matrix of the 60 measurements (30 lines by two replicates per line). We ran the model for a burn-in period of 2,000,000 samples then retained 1,000 posterior samples at a thinning interval of 100. Each parameter point estimate was then constructed from the mean of the corresponding distribution of 1,000 posterior samples.

### Convergence diagnostics

To determine whether the model had converged we considered autocorrelation across posterior samples for each estimated parameter. Approximately 1.5% of the trait loading estimates were associated with posterior autocorrelations exceeding 0.1. However, we were satisfied that the model had achieved a sufficient degree of convergence to proceed with interpretation. In particular, posterior means for PCF trait loadings and PCF scores were largely unchanged between sets of posterior samples taken after a burn-in period of 1,000,000 or 2,000,000 samples, as well as a previous implementation of the model in MATLAB (not shown).

### Significance testing

To determine statistical significance of PCF trait loadings and of PCF heritability we used the local false sign rate (LFSR) approach with an average error rate of 1%, as described in Hine *et al*. (2018). Briefly, LFSR is the probability of assigning the incorrect sign to an estimate. For PCF trait loadings, we assigned LFSR values as the proportion of posterior samples of the trait loading that were equal to zero, or on the other side of zero from the median of the posterior samples. We computed an average error rate, *s*, which is analogous to Storey’s q-value (Storey 2003; Stephens 2017), and considered a PCF to be statistically supported when loadings for a minimum of two traits had s <0.005 (2-tailed test as loadings can be positive or negative; false positive error rate of 1%). In *D. serrata*, all 59 estimated PCFs were statistically supported, ranging from 2 to 771 significant trait loadings (Fig. S1, Table S1), and collectively explained 59% of the phenotypic variance. Individual gene expression traits were significantly influenced by up to eight PCFs (Table S1). In *D. melanogaster*, 47 of the 59 PCFs were statistically supported, ranged in number of significant trait loadings from 2 to 549 (Fig. S1, Table S2), and collectively explained 39% of the total phenotypic variance. Individual *D. melanogaster* gene expression traits were significantly influenced by up to five PCFs (Table S2).

For PCF heritability, we assigned LFSR as the proportion of posterior samples equal to zero, and accepted a PCF as significantly heritable when s < 0.01 (i.e., 1% false positive rate for a 1-tailed test as variances cannot be negative), but additional requirements were imposed due to the potential for spurious associations in this data (randomisation analysis, detailed below). In *D. serrata* and *D. melanogaster*, respectively, 32 and 22 PCFs were significantly heritable, meeting the additional criteria from the randomisation analysis (Fig. S2). In *D. serrata*, heritability estimates for these heritable common factors (HCFs) ranged from 0.33 to 0.99 (Figures S1, S3), with individual HCFs significantly influencing between 4 and 483 traits, while individual traits were significantly influenced by to up to six HCFs (Table S1). In *D. melanogaster*, HCF heritability ranged from 0.30 to 0.99 (Figures S1, S4), significant trait loadings ranged in number from 5 to 549, and individual traits were influenced by up to three HCFs (Table S2).

### Identification of outlying lines

Other studies investigating the distributions of relationship between allelic effect size and frequency have defined effect size quantitatively (e.g., as the rank: Kremling *et al*. 2018), or qualitatively by defining outlier genetic variants based on a threshold z-score (reviewed in Table 1 of Richter *et al*. 2019). Here, to ensure robust detection of lines with unusual trait expression (outlier lines), we quantified effect size in units of trait-specific interquartile ranges from the trait-specific median. As outlined earlier, the BSFG analyses were conducted on variance-standardised data (i.e., on the trait-specific z-scores), which allowed us to interpret the square of the resulting PCF trait loadings as approximately the proportion of variance in the trait that can be explained by that PCF. However, we observed that when multiple visibly extreme values were present for a given trait, the inflation of variance due to the most extreme value sometimes resulted in z-scores of less extreme (but still visibly outlying) values being less than 2 SD from the mean, and therefore undetected as outliers.

Therefore, to ensure that we were capturing all extreme values in our identification of outliers, interrogation of the distributions of individual gene expression traits and PCF scores were conducted on the scale of interquartile range deviations (IQR). These characteristics of our data are consistent with general trends for gene expression data: while most of 100 GEO or NCBI gene expression datasets from 20 species were normally distributed, many followed a t-distribution, which is similar to the normal distribution but with heavier tails (i.e., extreme values occur more frequently) (Liu *et al*. 2019).

We identified outlier lines for all individual gene expression trait measurements (Table S3), and for PCF scores (Figures S3 and S4), as lines that satisfied each of three criteria. First, the mean of the two replicate values per line was greater than 3 IQR from the median of the 30 line means for that trait. Second, both replicate values per line were greater than 3 IQR from the median of the 60 values for that trait. Third, line replicate values were extreme deviates in a consistent direction (i.e., both replicates of the line fell on the same side of the median). The second and third criteria ensured that we were uncovering extreme genetic variants, not extreme values generated by spurious micro-environmental or technical effects on some individuals.

### Determination of spurious patterns through data randomisations

For datasets such as ours, where the number of variables (3,385) is far greater than the number of objects measured per variable (two replicates per 30 lines), random correlations among variables are expected to be common (e.g., Johnstone 2001). Therefore, to help assess the biological relevance of the HCFs, we implemented the BSFG model on 100 randomised versions of each dataset. The randomisation preserved the phenotypic and genetic variances of each trait individually, but disrupted the biological covariance among traits, resulting in an exact measure of the effect of sampling error on covariance, given the size of our experiment. To achieve this, we shuffled the data independently for each trait, randomly re-assigning the pair of replicate measures per line.

Across the 100 randomised datasets for each species, 46% of PCFs for *D. serrata* and 52% for *D. melanogaster* were significantly heritable (Figure S2), a surprising outcome given that in our previous application of the BSFG model to *D. serrata* data for the same traits measured in mutation accumulation (MA) lines, ≪1% of PCFs estimated in the randomisation analysis were significantly heritable (Hine *et al*. 2018). The higher number of spuriously heritable PCFs observed here may simply reflect differences in sampling error: the reduced number of lines in the current datasets (30 vs 41 in the MA lines), in combination with greater standing genetic than mutational variance, may have resulted in more spurious association among traits in the randomised standing genetic variance datasets. This effect would be exacerbated by the presence of extreme outliers (Results; Table S3), which were not observed in the analysis of the MA lines (Hine *et al*. 2018).

We further investigated the randomised data by characterising the PCFs estimated for each randomised dataset in terms of: i) the total number of significant trait loadings; ii) how many of those significantly loading traits had at least one line with an extreme value (“outlying-line traits”, defined above) and; iii) whether the PCF was significantly heritable. For both species, we observed a distinct relationship between these variables in the randomised datasets. The higher the proportion of outlying-line traits for a given PCF, the more likely the PCF was to be significantly (but spuriously) heritable (Fig. S2). These observations suggest that the frequent sampling of extreme values of multiple traits when assembling a random “line” via our shuffling approach (keeping both replicate measures of a line together) was not solely due to small number of lines but also (and perhaps more importantly) due to the true distribution of genetic effects per trait.

Given these observations, we used the spurious associations in the randomised datasets as a baseline against which to assess biological signal in the observed data. First, we considered observed data HCFs with a greater number of significantly loading traits than were recorded for any randomised dataset HCF to reflect biological signal, not sampling error. For the *D. serrata* data, most (29 of 35) HCFs had more significantly loading traits than any of observed for any of the spurious HCF observed in the 100 randomised datasets, but there was more overlap in significantly loading trait counts of randomised and observed data in *D. melanogaster* (Tables S1–2; Fig. S2). Where the range of significantly loading traits overlapped between observed and randomised dataset HCFs, we identified matched subsets of randomised data HCFs for which the number of significant trait loadings exactly matched that of a HCF in the observed dataset. We then compared the number of these significantly loading traits that were associated with an outlying line in the observed and the paired subset of randomised data HCFs (Fig. S2). If the number of outlying-line traits of the observed HCF fell outside the 95% range of the size-matched randomised HCFs, we deemed the observed HCF to be associated with a level of covariance that was distinguishable from sampling error and retained it for further investigation. The remaining HCFs, not distinguishable from spurious sampling association, were not considered further.

### Imposed orientation of PCF trait loadings and PCF scores

The orientation of the estimated PCF trait loadings is arbitrary, such that the direction could be reversed for any given PCF by multiplying its trait loadings by −1. Changing the direction of a PCF in this way will also result in the reversal in sign of the corresponding PCF scores. To aid interpretation, we imposed directionality on PCFs by multiplying the trait loadings and scores by the sign of the mean trait loading (i.e., by +1 or −1, depending on whether the average loading was positive or negative). This allowed us to investigate whether extreme variants were more likely to be associated with an overall increase or decrease in gene expression across the variance-standardised traits significantly affected by the latent factor.

### Investigating physical linkage of expressed genes

The BSFG approach implicates a latent genetic factor as causing the observed co-expression of genes that are significantly associated with the same HCF. Other studies have identified pleiotropic loci that affect the expression of large numbers of genes (e.g., Wang *et al*. 2010; Lukowski *et al*. 2017), and as such it is plausible that HCFs capture latent genetic factors that are pleiotropic loci. However, the experimental data analysed here cannot directly determine the genetic variant(s) responsible for the co-expression revealed by the BSFG, and thus whether pleiotropy is responsible. However, we can further address whether co-expressed genes were co-localised in the genome (*i.e.,* in linkage disequilibrium).

Allen *et al*. (2013) mapped 95% of all ESTs on the microarray analysed here to *D. melanogaster* chromosomes. Subsequent publication of the *D. serrata* genome assembly (available on NCBI: BioProject: PRJNA355616) (Allen *et al*. 2017) was consistent with the previous inference of strong gene location conservation between *D. serrata* and *D. melanogaster* (Stocker *et al*. 2012). A more recent scaffolding using DovetailTM Hi-C technology (Dovetail Genomics) greatly improved the contiguity of the assembly from an N50 of just under 1 Mbp (Allen *et al*. 2017) to an N50 of 30.3Mbp (S. Allen pers comm). Chromosome locations of the constructed scaffolds were determined based on 78 physical and linkage markers with known chromosome location (as identified by Stocker *et al*. 2012).

We queried the sequences of the 3,385 ESTs against the scaffolded *D. serrata* reference using the default settings in BLASTN (version 2.2.27+) (Altschul *et al*. 1990). This resulted in 94% of the ESTs mapping to the chromosomes X, 2L, 2R, 3L and 3R, which were, respectively, associated with 446, 545, 683, 688, and 817 ESTs. ESTs that did not successfully align with the reference genome were categorised as “other”, and are likely to have been derived from genes on chromosomes 4 or Y. We assumed the frequencies with which ESTS mapped to chromosomes were representative of an underlying multinomial distribution and used chi-square tests to determine whether subsets of ESTs corresponding to each HCF were distributed non-randomly across chromosomes.

To investigate the potential role of linkage disequilibrium in the identification of *D. melanogaster* HCFs, we focused specifically on the inversions that have been identified in the *Drosophila melanogaster* DRGP lines (Huang *et al*. 2014). Inversion genotypes (downloaded from the DRGP website: http://dgrp2.gnets.ncsu.edu/data.html) were available for 29 of the 30 lines analysed here. Across these 29 lines, five known inversions were segregating, present as one or two copies across 11 lines (including one line carrying two of the five inversions). Two of these five inversions, *In(2L)t* and *In(3R)Mo*, are associated with variation in the expression levels of hundreds of genes (Lavington and Kern 2017). Here, we were particularly interested in whether large changes in gene expression (i.e., HCFs with outliers) could be attributed to the presence of any of these inversions.

### Investigating differentiation in genetic roles of different classes of HCF

We conducted functional enrichment analyses on the subsets of genes represented by HCFs. For *D. melanogaster*, we used FlyBase Gene IDs corresponding to the Affymetrix probe set IDs as listed on the Gene Expression Omnibus (accession number GPL1322). Of the 3,385 probe sets analysed, 3,119 were associated with a single FlyBase Gene ID (110 had none, while the 156 probe sets associated with multiple FlyBase Gene IDs were also omitted). For *D. serrata*, we used FlyBase gene IDs corresponding to orthologs in *D. melanogaster* of the *D. serrata* transcript-level coding sequence, matched to *D. serrata* ESTs using BLASTN (default settings, version 2.2.27+) (Altschul *et al*. 1990). Orthologs were assigned using OrthoDB (Kriventseva *et al*. 2018). Transcript-level coding sequence was extracted from the annotation GFF file at NCBI Drosophila serrata Annotation Release 100 (https://www.ncbi.nlm.nih.gov/gene/annotation_euk/Drosophila_serrata/100/) using GffRead (Pertea and Pertea 2020). At least one ortholog was identified for 2,486 of the 3,385 *D. serrata* ESTs, including five ESTs with multiple orthologs and 90 orthologs that were represented by between 2 and 4 ESTs, resulting in 2,154 unique *D. melanogaster* orthologs. Enrichment analyses were conducted using the *gost* function in the *gprofiler2* R package (Raudvere *et al*. 2019) using the false discovery rate method of multiple testing correction with a threshold of 0.05. Queries were made against annotated genes from custom backgrounds corresponding to the 3,119 (*D. melanogaster*) or 2,154 (*D. serrata*) FlyBase Gene IDs. Limiting the background to only the genes that could have been in the gene set of interest reduces the probability of detecting significant enrichment where there is none (Timmons *et al*. 2015). We implemented semantic similarity analysis on any identified enriched terms for each HCF using the R package GOSemSim (Yu *et al*. 2010).

### Data availability

The full *Drosophila serrata* dataset is available at the Gene Expression Omnibus (GEO) database under accession number GSE45801. Due to a server fail, the exact version of the *Drosophila melanogaster* data originally reported in Ayroles *et al*. (2009) is no longer available. We have therefore used the data as summarized for the original demonstration of the BSFG presented by Runcie and Mukherjee (2013). The subsets of 3,385 expression traits measured in two replicate pools of male RNA from each of 30 lines analyzed here, for both species, along with the R code to implement the BSFG analyses and generate the parameters considered in this manuscript, can be downloaded from UQ eSpace (https://doi.org/10.48610/a3c5652).

## Results

Investigating the 32 *D. serrata* and 22 *D. melanogaster* Phenotypic Common Factors (PCFs) putatively caused by latent genetic factors (i.e., the Heritable PCFs, HCFs), we identified two distinct patterns that were present in both datasets. First, for 7 (22%) of the HCFs in *D. serrata* and 11 (50%) of the HCFs in *D. melanogaster*, there were no outlying lines (Fig. S3–S4; e.g., *D. serrata* HCF 8, Fig. 1 top left). These HCFs exhibit the characteristics expected with many small-effect contributions to genetic variation. Second, for the other 25 (78%) and 11 (50%) HCFs in *D. serrata* and *D. melanogaster*, respectively, there was at least one outlying line (Fig. S3–S4; Fig. 1 bottom left), a pattern consistent with expectations when genetic variance is due to rare, large effect variants. In *D. serrata*, there were 40 extreme values (4.2%) across the 32 HCFs and 30 lines. Most of the *D. serrata* HCFs with extreme values had only one outlying line (14 HCFs), but some had two or three outlying values (seven and four HCFs, respectively) (Fig. S3). Every *D. serrata* line was extreme for at least one HCF, while eight lines were extreme for two and one line was extreme for three HCFs (Fig. S3). In *D. melanogaster*, there were 11 extreme values (1.7%) across the 660 (30 lines × 22 HCFs) HCF values, with at most a single extreme value for any given line or HCF (Fig. S4).

**Figure 1.**
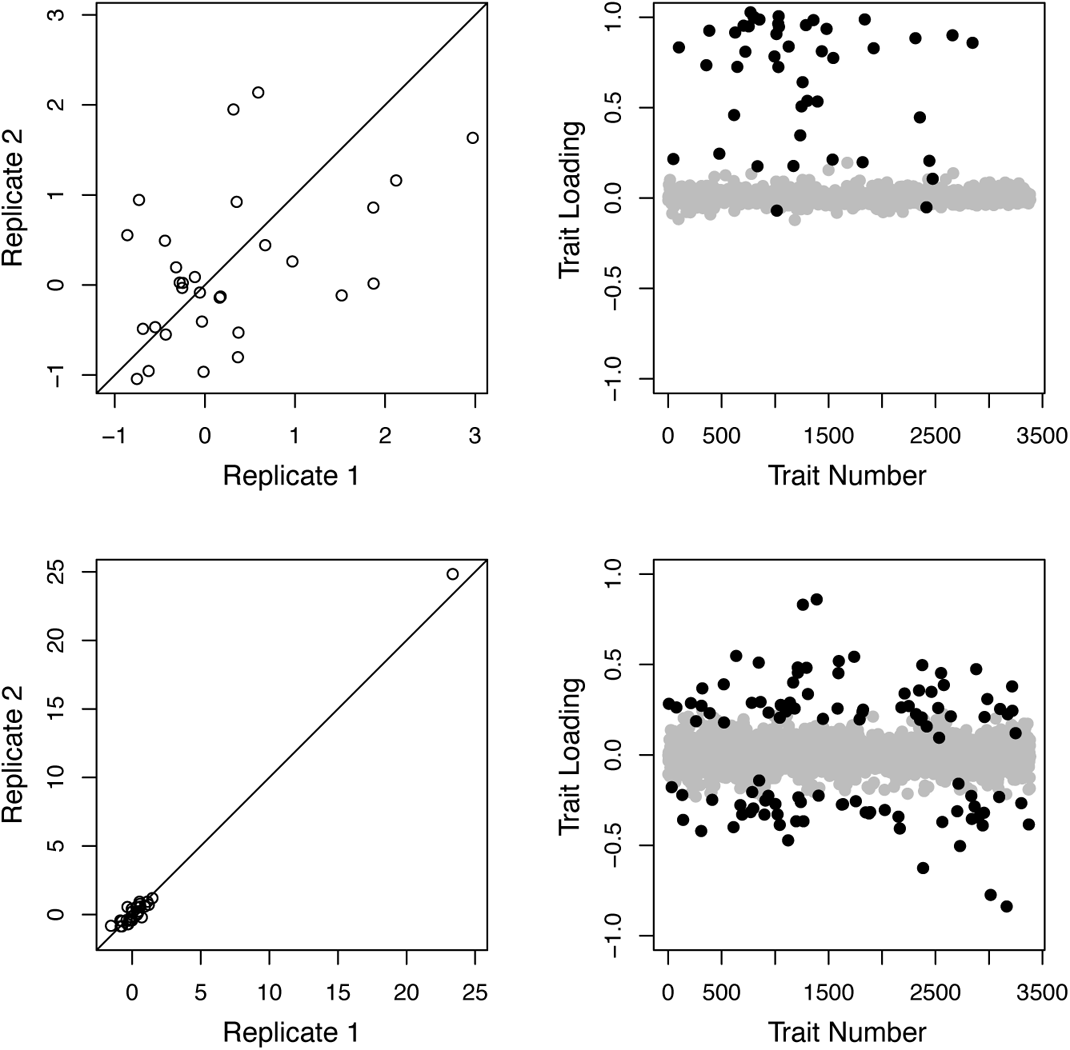
Example heritable PCFs from the *D. serrata* dataset, illustrating patterns commonly associated with HCFs with or without outlier lines. The distribution of estimated PCF scores for HCF 26 (no outlier lines; top left) and HCF 11 (one outlier line; bottom left) from the *D. serrata* dataset, on the scale of interquartile ranges from the median. The 30 points on each panel in the left column represent the two replicate deviations per line. The corresponding distributions of trait loadings for these two HCFs are also illustrated (right column; black (grey) circles depict significant (non-significant) trait loadings), where traits were arbitrarily assigned a numerical identifier prior to analyses.

The presence or absence of outlier lines for an HCF was associated with differences in the nature of the HCF. In both species, HCFs with outliers exhibited higher heritability (Fig. 2, top row) and less bias in the direction of significant loadings (Fig. 2, second row). Specifically, for HCFs with no outliers, most significantly loading traits were influenced by the latent HCF in the same direction, corresponding to an axis of variation contrasting lines that upregulated this set of genes with lines that downregulated expression of all these genes, relative to the population mean expression levels (e.g., Fig. 1 top right; Fig. S5–S6). In contrast, HCFs with outliers exhibited a more symmetrical distribution of loadings (e.g., Fig. 1 bottom right; Fig. S5–S6). The number of traits significantly influenced by an HCF also significantly differed, but not consistently between the species. In *D. serrata*, HCFs with outliers had more significant loadings (Fig. 2, left panel, 3^rd^ row), while in *D. melanogaster* they had fewer (Fig. 2 right panel, 3^rd^ row). The average magnitude of significant trait loadings did not differ significantly between HCFs with versus without outliers for either species (Fig. 2 4^th^ row), although in *D. serrata*, loadings on HCF without outliers were noticeably at the lower end of the range relative to HCFs with outliers (Fig. 2 left panel, 4^th^ row). Thus, where a trait was significantly influenced by an HCF, the proportion of phenotypic variance in the trait arising from that HCF (approximated as the square of the trait loading) was, on average, not significantly different for HCFs with vs without outliers.

**Figure 2.**
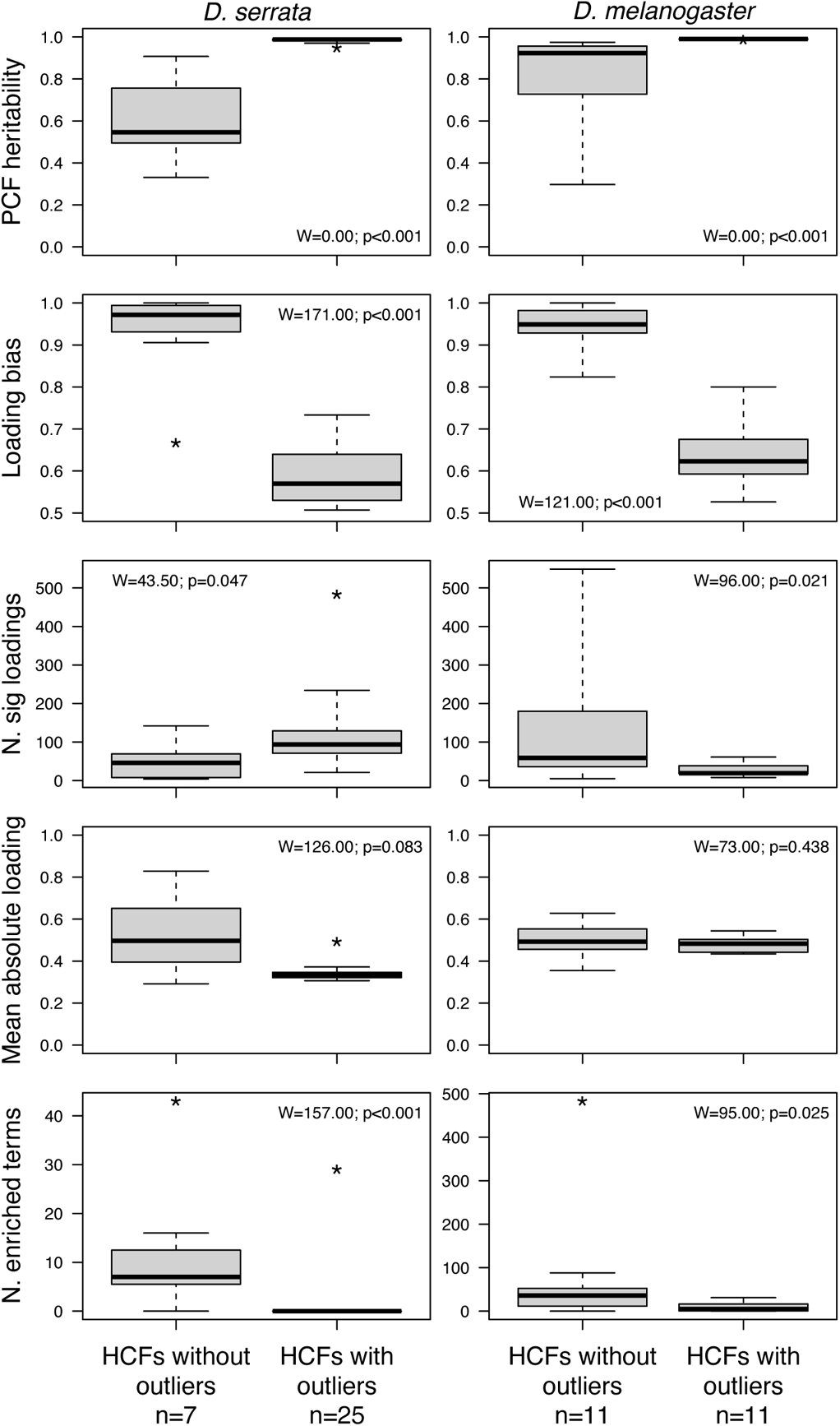
Comparison of characteristics of heritable PCFs with and without outliers. Bold line, box and whiskers represent the median, 1.5 interquartile ranges (IQR) and 3 IQR, respectively. Values exceeding 3 IQR are indicated with an asterisk. We compared each characteristic between the two types of heritable PCF (outliers absent or present) using the Wilcoxon Rank Sum Test (results shown within each panel).

We investigated whether outlier values for an HCF emerged as a simple consequence of lines being extreme for the individual traits influenced by the putative latent genetic factor. First, we determined that some individual gene expression traits were characterised by extreme values (outlier lines): of the 3385 traits measured, 3.9% (*D. serrata*) or 6.7% (*D. melanogaster*) had at least one line that was >3 IQR from the median (Table S3), with some lines deviating up to 18 (*D. serrata*) or 19 (*D. melanogaster*) IQR from the median. Next, we considered whether the presence of extreme values for individual traits predicted the presence of extreme values for the HCF(s) to which the trait contributed. In *D. serrata*, all individual expression traits with outlier line(s) were also associated with HCFs with outlier lines (eight traits, which significantly loaded onto more than one HCF, were also associated with a HCF without any outlier lines: Table S3). Similarly for *D. melanogaster,* traits with outliers were disproportionately associated with HCFs with outliers, although the relationship was not as definitive: 61.4% of traits with outliers contributed to HCF with an outlier, while 20.3% contributed to HCF without an outlier line, and 18.3% were significantly associated with both types of HCF (Table S3).

We further dissected this general pattern by explicitly asking whether individual lines (genotypes) that were extreme for a given HCF were a consequence of having outlier values for the individual traits that were significantly influenced by that HCF (Fig. 3). Overall, in both species, there was a significant positive association between a line’s HCF value and the proportion of individual traits for which it was also an outlier (Fig. 3). Individual lines that were not outliers on a given HCF (grey points in Fig. 3) also tended not to have extreme values of the associated individual traits (i.e., have a value of 0 on the y-axis in Fig. 3A, B). However, some of these HCF non-outlier lines did have extreme values for the associated individual traits, with lines > 3 IQR for up to 10% (*D. serrata*) or 20% (*D. melanogaster*) of individual traits. Thus, covarying extreme values for some traits did not inevitably result in extreme values for the line on the corresponding HCF, but lines that were outliers for a given HCF did nonetheless tend to have higher proportions of contributing traits that were outliers (Fig. 3). For *D. serrata*, some line(s) had extreme values for an HCF despite being an outlier line for only a small proportion (<10%) of the associated individual traits (bottom right of Fig. 3A). Notably, outlier lines for the HCFs that had only one outlier line (blue points in Fig. 3; 14 HCF in *D. serrata*, and all 11 HCF with outliers in *D. melanogaster*) tended to have a higher proportion of individual trait outliers than HCFs with multiple outliers (red or black points in Fig. 3).

**Figure 3.**
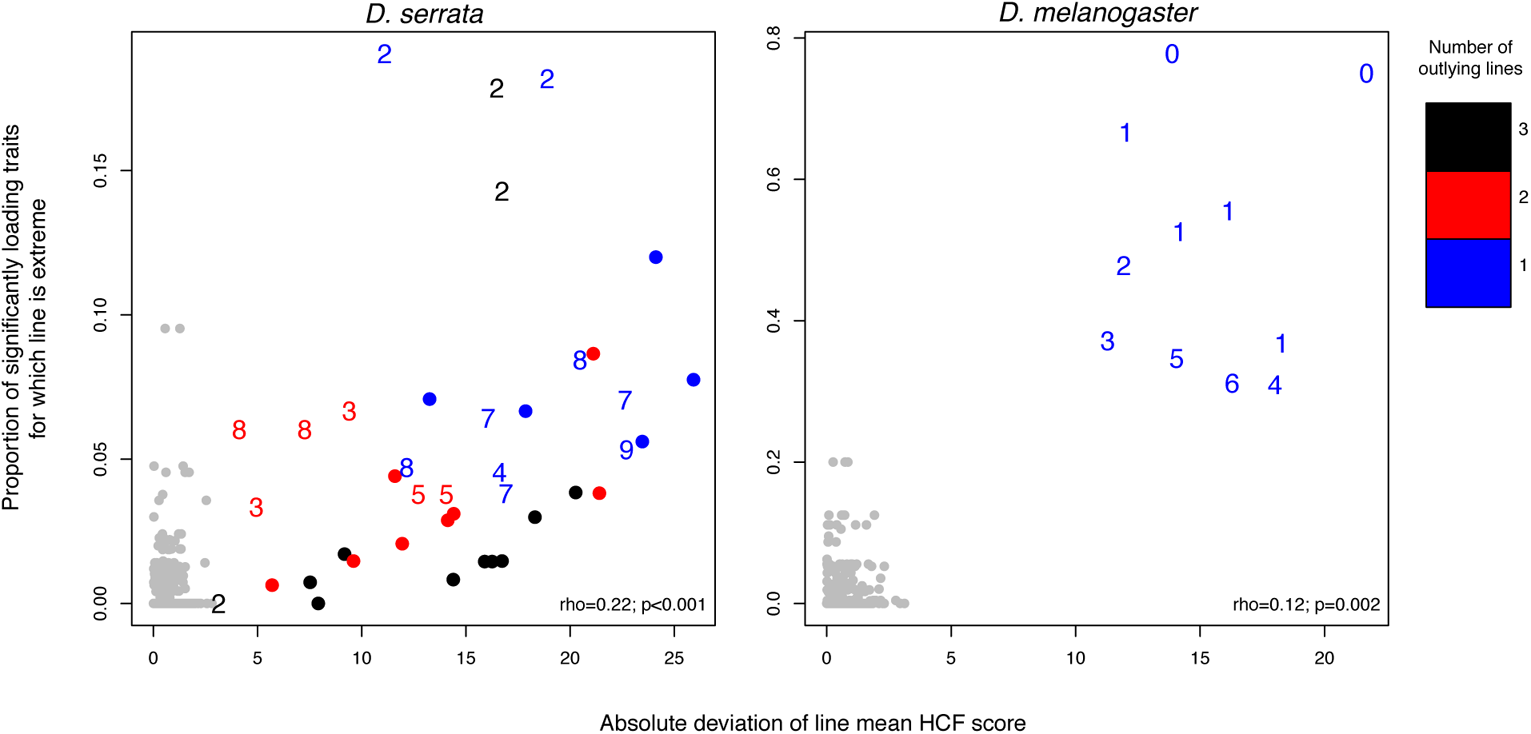
Association between size of the deviation of a HCF score and the frequency of extreme gene expression values for individual contributing traits. For each HCF (32 observations in *D. serrata* and 22 in *D. melanogaster*), we plotted each of the 30 line-mean deviations (x-axis) and the proportion of individual traits that loaded significantly to that HCF for which the line was also an outlier (y-axis; note the different scale for each species). For each HCF, lines that were not outliers (<3 on x-axis) are shown as grey points. For lines that were outliers on an HCF, plot symbols (numbers) indicate the number of significant trait loadings, in steps of 10, from “0” indicating an HCF with <10 loadings through to a dot for >100 loadings (Tables S1–2). Colours indicate the number of outlier lines for that HCF (see figure for key). For example, a red “3” indicates an HCF with two extreme line-mean values (red) and between 30 and 39 significant loadings (3). Spearman’s correlation between line mean HCF score and proportion of individual traits outliers are shown within each panel.

Having observed that HCFs with outliers had significantly less bias in the direction of their individual trait loadings (Fig. 2, 2^nd^ row), we further investigated the distribution of the extreme values on HCFs with outliers (25 HFC in *D. serrata* and 11 in *D. melanogaster*), and the individual expression traits they were predicted to affect. Note that the imposed PCF directionality (see *Methods*) allowed us to infer positive (negative) extreme PCF scores to indicate an overall increase (decrease) in variance-standardised gene expression when summed across the traits significantly associated with the PCF. In *D. serrata*, 33 of the 40 outlier values across all 25 HCF deviated above the median, while only 7 deviated below (Fig. 4A; Fig. S3). This directional bias was less obvious for individual traits loading significantly on these HFCs, with many individual traits deviating >3 IQR below the median (Fig. 4A). In contrast, in *D. melanogaster*, extreme variants for HCFs were evenly distributed above versus below the median (Fig. 4B, Fig. S4), although the most extreme values were below the median, and individual traits were biased towards decreased expression (Fig. 4B).

**Figure 4.**
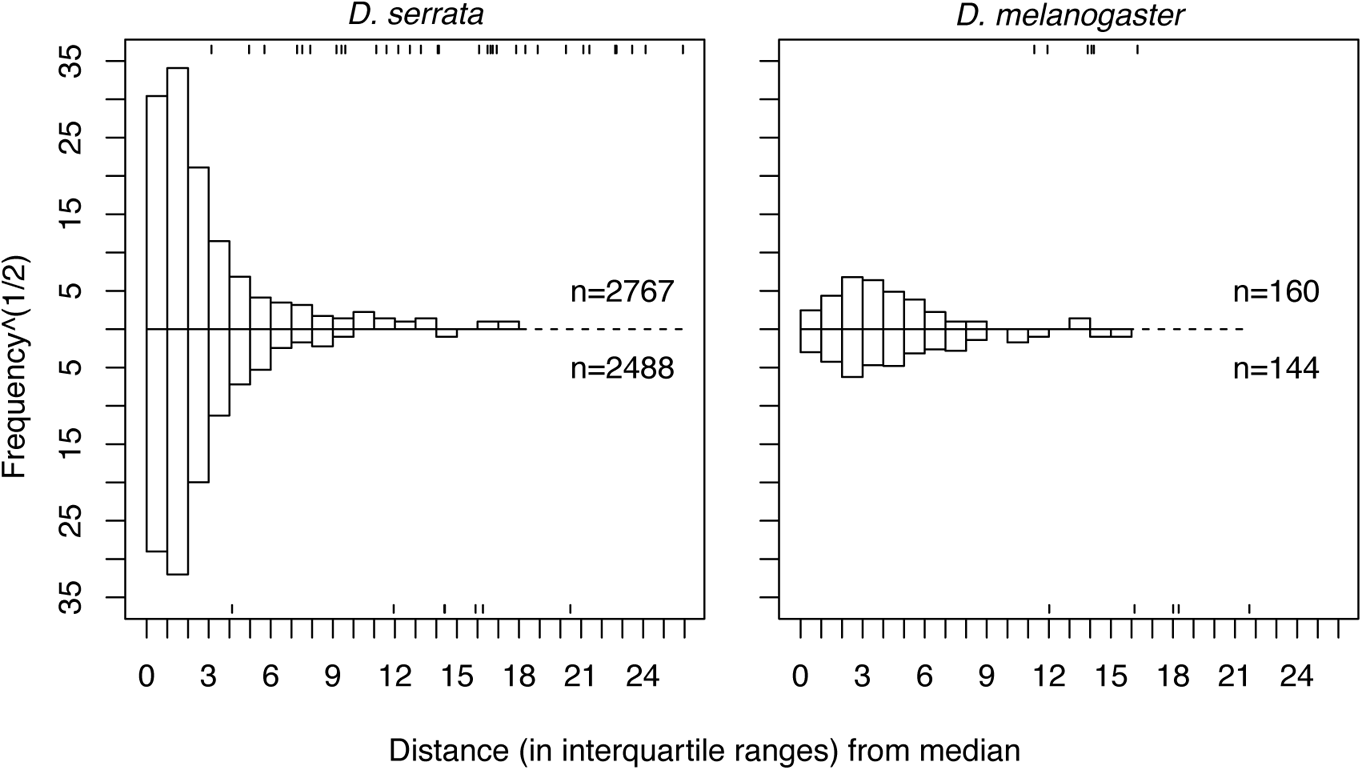
Directionality of extreme line-mean deviations for HCFs and their corresponding gene expression traits. Line-mean deviations of HCFs are shown as whiskers at the top and bottom of each panel, corresponding to positive and negative deviations, respectively. The reflected histograms show the square root counts of positive (above the dashed line) and negative (below the dashed line) line-mean deviations of traits associated with HCFs with extreme values, for the HCF-specific lines with extreme HCF values.

### Is coexpression on HCFs consistent with physical linkage?

In *D. serrata*, we determined whether genes influenced by the same latent genetic factor co-localised in the genome. Observed chromosome frequencies did deviate significantly from expected frequencies for seven of the 32 HCFs (Table S4). However, in only one case were all genes associated with a given HCF co-localised to the same chromosome (arm): 10 traits associated with HCF31 did not map to a major chromosome, suggesting they occur on Y or 4 (Table S4). Notably, HCFs for which gene co-localisation was statistically supported represent both those with extreme outlier lines (HCFs 2, 4, 20, 22 and 23) and those without outliers (HCFs 26 and 31) (Table S4).

In *D. melanogaster*, we investigated whether co-localisation of genes within an inversion could account for the large changes in gene expression in the lines with outlying HCF scores. However, there was no consistent association between outlier status and inversion karyotype for any of the HCFs with outliers (Table S5). For any given inversion and HCF combination, there were either no outlier lines with the inversion, or there was at least one representative of each class of line (with versus without an outlier) that had one or more copies of the inversion.

### Functional characteristics of HCF

In *D. serrata*, six of seven HCFs without outliers were significantly enriched for terms in one or more of the GO categories (BP - biological process, CC - cellular component and MF - molecular function; Table S6). HCF29 was associated with the largest number of terms, which broadly related to development. Of the 25 HCFs with outliers, only HCF22 was significantly enriched, with 29 BP terms relating to meiosis, and the detection of stimulus or taxis. In *D. melanogaster*, of the 11 HCFs without outliers, 10 were significantly enriched for terms in at least one of the GO categories (Table S6). Eight of the 11 HCFs with outliers were significantly enriched for terms from one or more of the BP or MF categories, while terms from the CC category were notably absent; the overall number of enriched terms was substantially lower for HCFs with outliers than those without (Table S6; Fig. 2, bottom row). Given the larger number of trait loadings for *D. serrata* HCFs with versus without outliers (Fig. 2), we considered whether this could have caused the pattern of differential enrichment observed. However, we note the numbers of genes in the focal and background lists are key parameters of the hypergeometric test, accounting for differences in gene number among focal lists. Further, despite the overall mean difference in gene number, many HCFs with outliers had comparable numbers of genes in the analyses. For example, HCFs 27-29 (without outliers) were associated with the largest number of enriched terms for *D. serrata* and had gene sets ranging in size from 44 to 123 genes (Table S6). HCFs 4-16, 18 and 19 (with outliers) all had gene set sizes within this range but were not enriched for any terms in the GO categories (Table S6).

## Discussion

The evolutionary dynamics of quantitative genetic variation remain poorly understood, with many competing models, none of which fully reconcile empirical observations on mutation, selection and standing genetic variation (Walsh and Lynch 2018). Direct tests of the predictions of different models require knowledge of the effective population size, the number of loci affecting traits, their mutation rate, and the variance in effects of those mutations, all parameters that are difficult to accurately estimate (Walsh and Lynch 2018). Furthermore, while pleiotropy has been implicated as playing an important role in determining the magnitude of genetic variance that can be maintained in the presence of selection (Johnson and Barton 2005; Walsh and Lynch 2018), we lack the empirical assessments of high-dimensional data necessary to consider model predictions. Here, we interrogated the multivariate distribution of genetic variance in quantitative phenotypes to characterise patterns of shared variation among traits. Below, we consider the implications of our observations for the maintenance of genetic variance in natural populations.

For individual traits, rare direct evidence (Hodgins-Davis *et al*. 2015) and indirect evidence from genetic mapping studies in humans and model taxa (Battle *et al*. 2014; Bloom *et al*. 2019) support House of Cards (HoC) over Gaussian mutation models of the maintenance of genetic variance. Consistent with this emerging body of evidence, we detected lines (genotypes) with extreme values for individual gene expression traits in two species of *Drosophila*. Importantly, rare extreme variants were also present for multivariate combinations of these individual traits, suggesting that the HoC mutation model may also account for heritable variation of higher-dimension phenotypes. At least half of the multivariate traits influenced by a latent genetic factor (i.e., the heritable common factors, HCFs) had at least one extreme genotype (line), resulting in higher heritability of these multivariate traits. While multivariate (HCF) extreme values typically occurred for lines (genotypes) with a higher proportion of individual trait outliers, individual traits could be extreme without resulting in extreme values for HCF that they were influenced by, and conversely, extreme HCF values were observed when the latent factor did not cause extreme values of any individual trait (Fig. 3). Thus, the presence of rare, extreme values for multivariate expression traits could not be simply predicted from inspection of the individual trait distributions.

Gaussian and HoC models differ in predictions of the evolutionary dynamics underpinning standing genetic variation in equilibrium populations, where under the HoC model, genetic variation is determined by strong selection against rare, large effect mutation (Kingman 1978; Turelli 1984; Burger *et al*. 1989; Walsh and Lynch 2018). Given several simplifying assumptions, we make some tentative observations concerning the strength of selection in the natural *Drosophila* populations. First, we presume that extreme trait values (including for HCFs) are caused by single loci, where for HCFs these loci have pleiotropic effects. *Trans*-regulatory factors can influence expression of large numbers of genes (Brem *et al*. 2002; Albert *et al*. 2018; Cesar *et al*. 2018), and generate extensive genetic co-variation of expression (Denver *et al*. 2005; Lukowski *et al*. 2017). Therefore, it is plausible that the observed patterns of covariation reflect trans-acting pleiotropic alleles. Chromosomal location of loci, or inheritance of chromosomal inversions, were not consistent with loci affected by the same latent factor (HCF) being in close physical linkage, but nonetheless, we cannot exclude chance sampling of extreme alleles at independent loci (and the resulting transient linkage) as causal latent factors for the HCF; we further consider the nature of the latent genetic factors below.

Given the presumption of a single (pleiotropic) locus, and the observation that, for most individual traits and HCFs, only one extreme line was observed, we infer that alleles with (homozygous) major effects were segregating in the base population at a frequency of *q* ≈ 1% (~1 in the ~120 genomes sampled to found the panel of 30 lines in these diploid species). Applying equation 2.13 of Falconer and Mackay (1996), 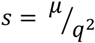, where *μ* is the genic mutation rate, predicts that the strength of selection required to maintain *q* at 1% was ~0.01 − 0.1, given *μ* of 10^−5^ − 10^−6^ (Lynch and Walsh 1998). If the true *q* is in fact lower (higher) than our sample suggests, *s* must be stronger (weaker) to maintain the allele frequency, assuming *μ* remains the same. Genomic mutation rates are heterogeneous across the genome (Nachman and Crowell 2000; Hodgkinson and Eyre-Walker 2011; Smith *et al*. 2018; Nesta *et al*. 2021), and mutations with larger effects might occur more rarely than smaller effect mutations (Mackay *et al*. 1992; Davies *et al*. 1999; Eyre-Walker and Keightley 2007; Heilbron *et al*. 2014; Mcguigan *et al*. 2014b; Kim *et al*. 2017); such variability, resulting in higher (lower) genic mutation rates would correspond to stronger (weaker) selection to maintain *q* ~ 1%. Finally, this prediction assumes that the large-effect alleles are recessive; if the observed outlier lines in fact carry alleles with additive or dominant effects, then the resultant increased visibility to selection means that selection 50-100 times weaker could maintain alleles at 1% frequency (Falconer and Mackay 1996, eqns 2.14 & 2.15). The predicted strength of selection (*s* ~ 0.01 − 0.1) detailed above is within the range estimated for new mutations affecting fitness traits, such as viability and fecundity, based on the ratio of mutational to genetic variance (~0.02: Houle *et al*. 1996), and consistent with the estimated average deleterious effect of heterozygous lethal genes in wild and laboratory *Drosophila* populations (~0.02: Crow and Temin 1964). The ratio of mutational to genetic variance was also used to estimate *s* for smaller sets (five) of gene expression traits in the *D. serrata* lines analysed here, inferring selection within the same range for 5-dimensional multivariate axes of expression (median *s* = 0.032), but weaker selection on the individual gene expression traits (median *s* = 0.005) (Mcguigan *et al*. 2014a). Nonetheless, while the strength of selection inferred to be required to maintain *q* ~ 0.1 is consistent with other evidence, we need further information on mutation rates specific to loci that have been characterised as large effect variants, and on the frequency spectrum of those alleles in natural populations, to gain further insight into the mutation-selection interaction underpinning extreme multivariate trait observations.

Puzzlingly, we observed that while enrichment analyses identified co-expression patterns associated with gene functions for HCFs without outliers, there was little evidence of common function among HCFs with outliers, particularly in *D. serrata*. The two classes of HCF also differed in the pattern of co-expression. HCF lacking outliers showing biased direction of influence on individual expression traits, corresponding to lines with relatively high versus low expression of all affected traits. This is consistent with a previous analysis of these *D. serrata* lines, where Blows *et al*. (2015) applied a matrix-building approach to estimate a single multivariate axis of genetic variation in expression of 8,750 expression traits. This co-expression axis was characterised by biased direction of trait loadings (indicating co-ordinated up-down regulation of many genes), and was enriched for multiple GO terms related to transcriptional regulation (Blows *et al*. 2015).

HCF with outliers, which were less likely to be enriched for a specific function, exhibited patterns among lines where expression was elevated for some traits, but depressed for others (i.e., unbiased direction of loadings). There are several possible non-exclusive explanations for these observations, which our data cannot further distinguish among. HCF with outliers might capture gene co-expression caused by a mixture of so-called horizontal (e.g., shared chromatin status) and vertical (e.g., shared transcription factors) processes. Van Dyke *et al*. (2021) recently demonstrated that genomic “hotspots”, containing QTL with *trans*-effects on multiple expression traits (eQTL) cause coordinated changes in expression of functionally unrelated genes via horizontal mechanisms. Second, the observed pattern (large phenotypic effect, apparently unrelated function) of HCF with outliers could reflect extreme values of pleiotropic effects. The distribution of effects of pleiotropic alleles across multiple traits is largely unknown, but might be expected to include alleles with very weak effects on some trait(s) (Hill and Zhang 2012; Paaby and Rockman 2013); the HCF outlier patterns observed in this study could reflect rare pleiotropic alleles, with effects in the extreme tails of the distribution of joint effects. Alleles with such extreme pleiotropic effects might be selectively eliminated, or subject to epistatic modification to limit the range of biological processes influenced. Thus, a third plausible explanation of these data is that epistatic effects on gene expression might generate large phenotypic outliers through sampling effects (chance segregation within a single line of an unusual combination of alleles across physically unlinked loci), or, as suggested by Mackay (2014), the presence of recent mutation (i.e., rare allele) for which the population has yet to evolve epistatic amelioration of effects.

Quantitative genetic theories of the maintenance of genetic variance that incorporate indirect (apparent) stabilising selection assume that mutations have directionally biased effects on fitness itself (decreasing it), but that the pleiotropic effects of those mutations on other traits are unbiased, equally frequently increasing as decreasing trait values (Barton 1990; Kondrashov and Turelli 1992; Johnson and Barton 2005). Empirical data to assess the assumption are sparse. For the relatively well-studied trait of size, studies in several taxa suggest that mutations typically decrease body size (Keightley and Ohnishi 1998; Lynch *et al*. 1998; Azevedo *et al*. 2002; Estes *et al*. 2005; Ostrow *et al*. 2007), with larger effect (Santiago *et al*. 1992) or more deleterious (Mcguigan and Blows 2013) mutations being particularly biased. Intensive study of mutational effects in *Saccharomyces cerevisiae* found that mutations more frequently increased than decreased expression of two of the ten genes studied, while a third gene exhibited the opposite bias, with more mutations decreasing expression (Hodgins-Davis *et al*. 2019). Thus, directional bias of mutational effects might be more prevalent than appreciated. Here, in *D. serrata*, for HCF with outliers (and for the traits influenced by these putative latent genetic factors), expression was biased upward; the question of emergent strong bias in the multivariate distribution of phenotypic effects warrants further investigation.

## Conclusions

Resolution of theory predicting the maintenance of genetic variation for quantitative phenotypes depends on better insight into the distributions of (pleiotropic) allelic effects on multiple traits and fitness. Widespread availability of data on many expression traits from the same individuals provides a particularly powerful system for investigating shared variation among high-dimensional phenotypes, while the intermediary causal nature of expression traits, connecting genotype to more complex traits (Li *et al*. 2017), adds to their appeal. Our results suggest that the growing evidence that genetic variance might be due predominately to relatively large effect variants, consistent with House of Cards mutation models, might extend to multivariate gene expression phenotypes. However, it also remains to be determined whether the simple genetic basis of the large latent factors inferred here could be peculiar to gene expression traits, where the potential for hierarchical control of gene regulation would lend itself to master regulators of expression. Therefore, further investigations of other types of traits are required to determine whether complex trait covariances are typically consistent with large effect mutation contributing strongly to standing genetic covariance.

## Acknowledgments

This work was supported by the Australian Research Council.

---

**Table S1.**
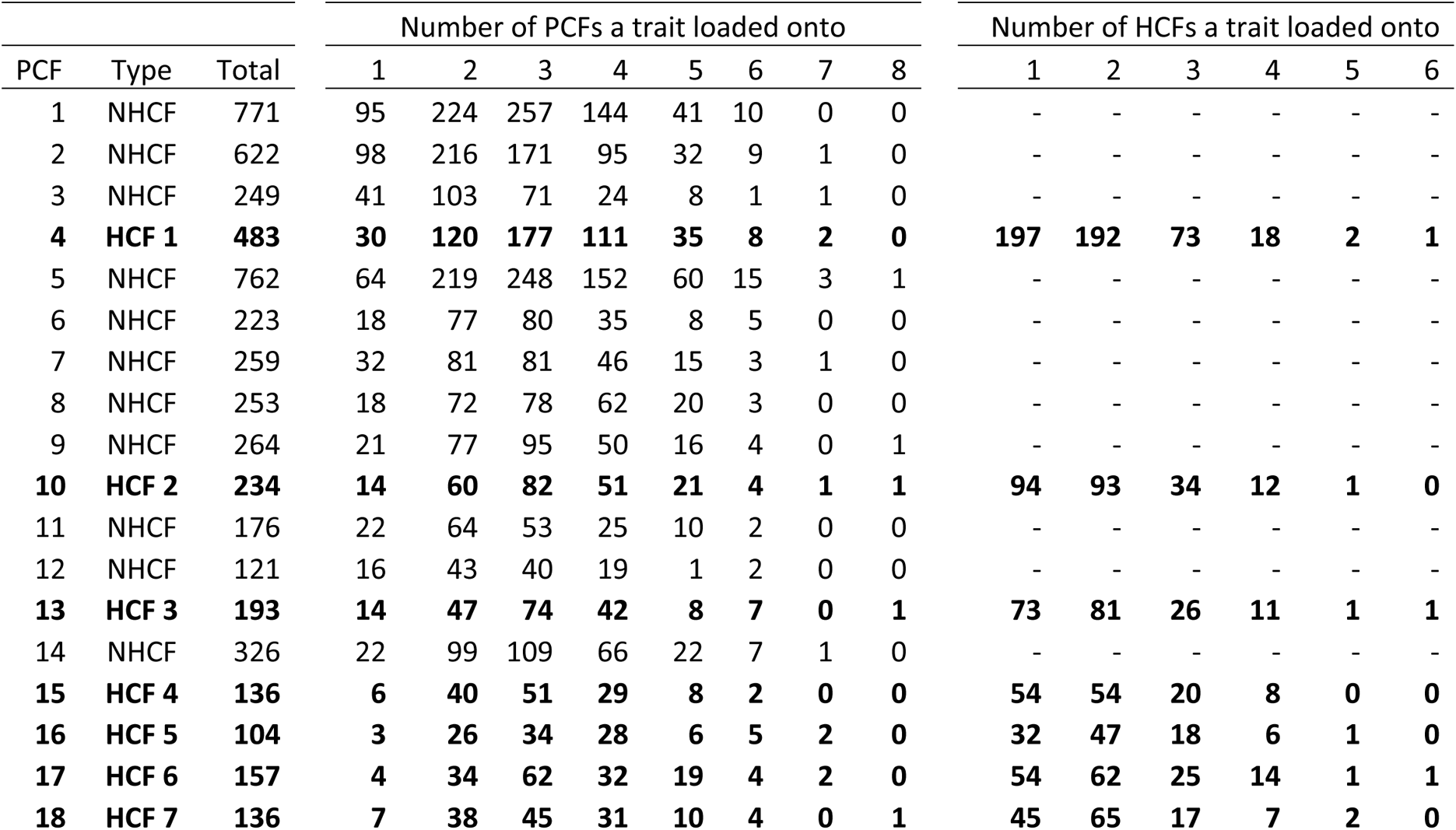
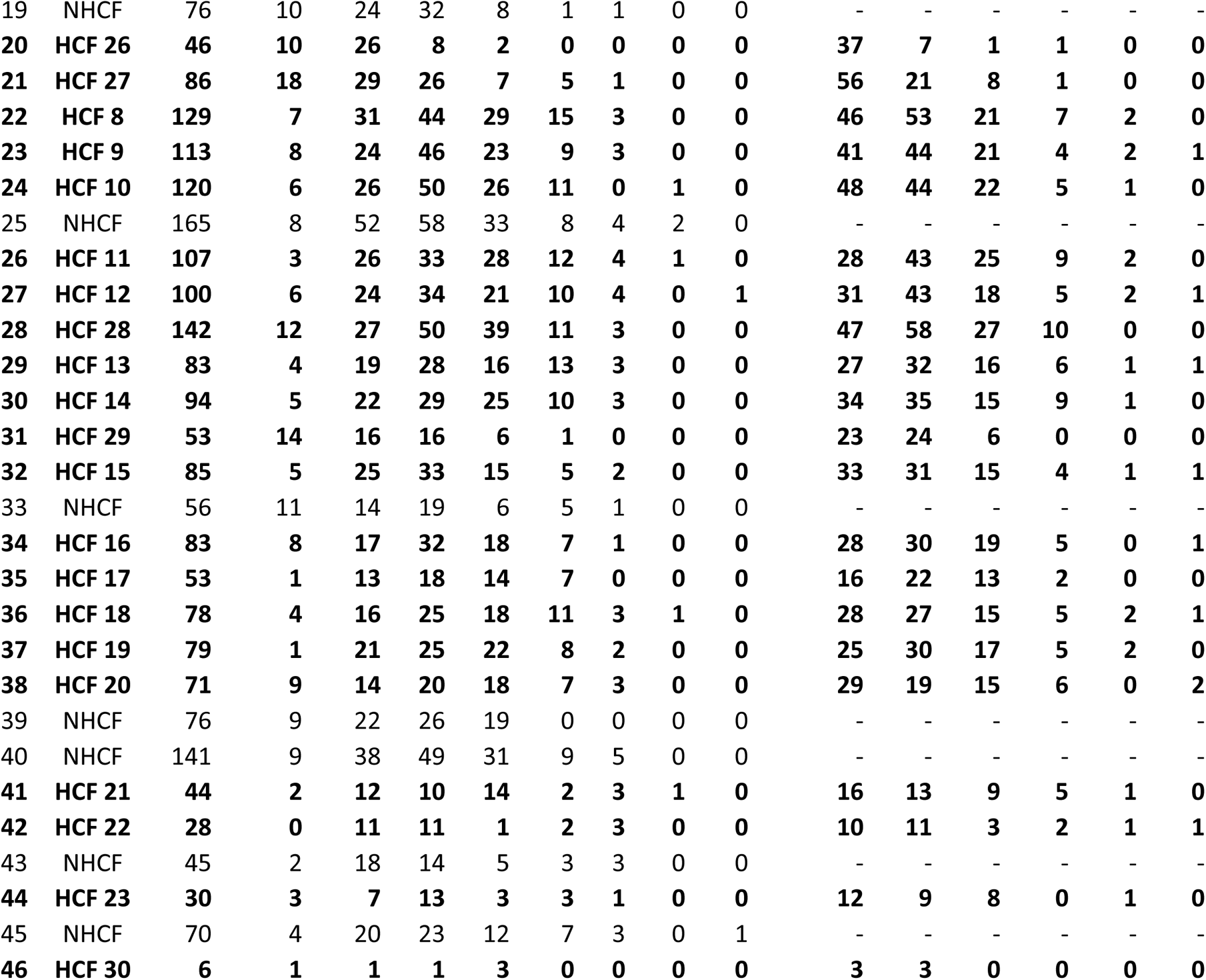
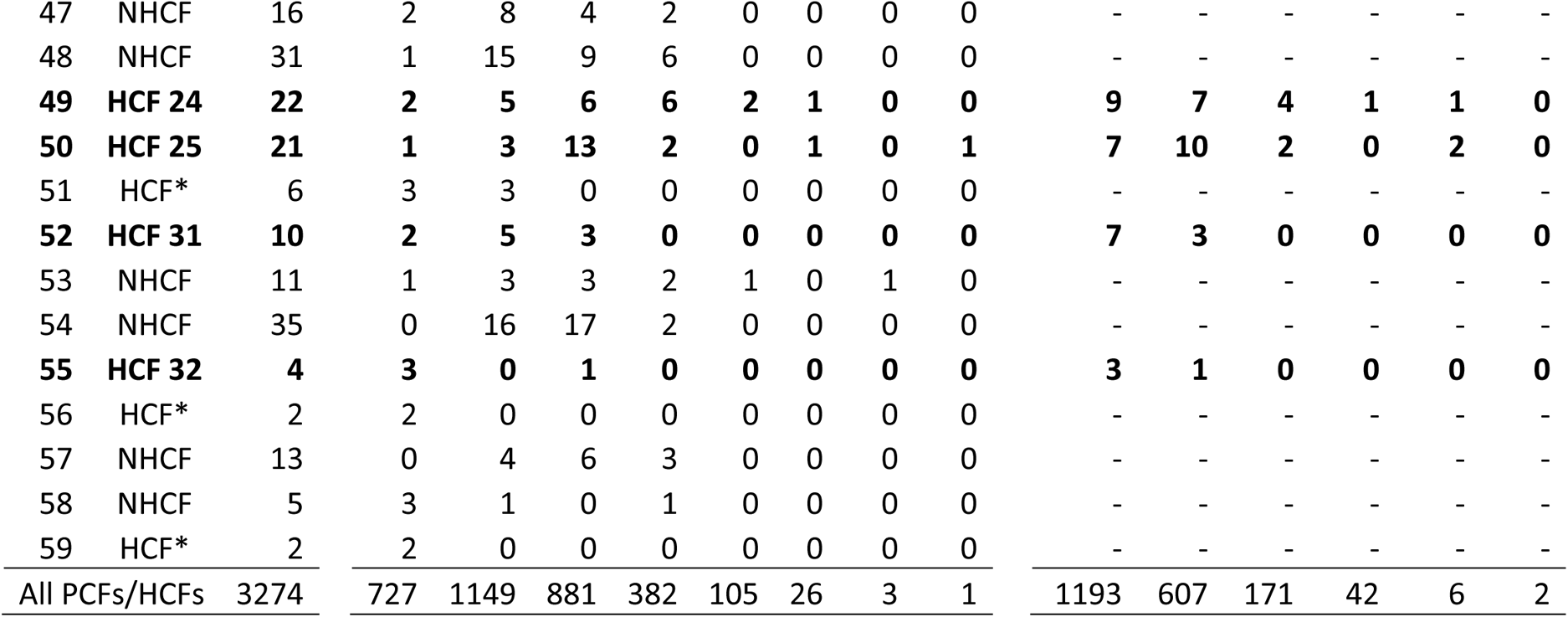
Number of traits associated with common factors estimated for the D. serrata gene expression dataset. For each phenotypic common factor (PCF), we report the total number of associated traits, then partition this number into how many PCFs these traits are associated with. PCFs are ordered by the magnitude of their predicted contribution to phenotypic variation. We further classify PCFs into three types (second column): PCFs that were not significantly heritable (NHCF), heritable common factors (HCF) and heritable common factors with trait profiles indistinguishable from that expected from sampling error (HCF*). HCFs (highlighted in bold) are numbered because they were subject to further analyses. For each of the HCF, we also partition the significant trait loadings by how many times a loading trait contributed to an HCF. The bottom row summarizes these numbers across all PCFs and HCFs.

**Table S2.** Number of traits associated with common factors estimated for the D. melanogaster gene expression dataset. For each phenotypic common factor (PCF), we report the total number of associated traits, then partition this number into how many PCFs these traits are associated with. PCFs are ordered by the magnitude of their predicted contribution to phenotypic variation. We further classify PCFs into three types (second column): PCFs that were not significantly heritable (NHCF), heritable common factors (HCF) and heritable common factors with trait profiles indistinguishable from that expected from sampling error (HCF*). HCFs (highlighted in bold) are numbered because they were subject to further analyses. For each of the HCF, we also partition the significant trait loadings of each HCF by how many times a trait contributed to an HCF. The bottom row summarizes these numbers across all PCFs and HCFs.

| PCF | PCF.type | Total | Number of PCFs a trait loaded onto |  |  |  |  | Number of HCFs a trait loaded onto |  |  |
| --- | --- | --- | --- | --- | --- | --- | --- | --- | --- | --- |
|  |  |  | 1 | 2 | 3 | 4 | 5 | 1 | 2 | 3 |
| **1** | **HCF 12** | **549** | **282** | **208** | **50** | **9** | **0** | **454** | **90** | **5** |
| 2 | NHCF | 410 | 215 | 141 | 48 | 6 | 0 | - | - | - |
| **3** | **HCF 13** | **125** | **63** | **53** | **8** | **1** | **0** | **104** | **20** | **1** |
| 4 | NHCF | 275 | 125 | 107 | 35 | 7 | 1 | - | - | - |
| **5** | **HCF 14** | **235** | **97** | **107** | **25** | **6** | **0** | **168** | **63** | **4** |
| 6 | NHCF | 438 | 155 | 189 | 83 | 11 | 0 | - | - | - |
| **7** | **HCF 15** | **236** | **99** | **89** | **41** | **7** | **0** | **167** | **61** | **8** |
| **8** | **HCF 16** | **65** | **32** | **24** | **8** | **1** | **0** | **50** | **14** | **1** |
| 9 | HCF* | 72 | 21 | 35 | 15 | 1 | 0 | - | - | - |
| **10** | **HCF 17** | **40** | **23** | **12** | **3** | **2** | **0** | **29** | **9** | **2** |
| **11** | **HCF 18** | **28** | **8** | **12** | **8** | **0** | **0** | **19** | **9** | **0** |
| **12** | **HCF 1** | **61** | **22** | **30** | **6** | **3** | **0** | **44** | **15** | **2** |
| **13** | **HCF 19** | **59** | **32** | **19** | **7** | **1** | **0** | **44** | **15** | **0** |
| **14** | **HCF 20** | **49** | **32** | **11** | **5** | **0** | **1** | **41** | **8** | **0** |
| **15** | **HCF 2** | **35** | **18** | **12** | **5** | **0** | **0** | **27** | **7** | **1** |
| **16** | **HCF 3** | **52** | **13** | **28** | **10** | **0** | **1** | **32** | **18** | **2** |
| **17** | **HCF\*** | **45** | **15** | **20** | **7** | **3** | **0** | **-** | **-** | **-** |
| 18 | HCF* | 37 | 13 | 12 | 12 | 0 | 0 | - | - | - |
| 19 | HCF 4 | 42 | 13 | 14 | 12 | 3 | 0 | 21 | 18 | 3 |
| 20 | HCF* | 40 | 15 | 17 | 8 | 0 | 0 | - | - | - |
| 21 | HCF* | 22 | 9 | 9 | 4 | 0 | 0 | - | - | - |
| 22 | HCF* | 36 | 11 | 18 | 6 | 1 | 0 | - | - | - |
| 23 | HCF* | 34 | 7 | 16 | 8 | 2 | 1 | - | - | - |
| 24 | HCF* | 33 | 10 | 15 | 7 | 1 | 0 | - | - | - |
| 25 | HCF 5 | 23 | 5 | 10 | 8 | 0 | 0 | 13 | 8 | 2 |
| 26 | HCF* | 25 | 8 | 8 | 7 | 1 | 1 | - | - | - |
| 27 | HCF* | 27 | 10 | 11 | 6 | 0 | 0 | - | - | - |
| 28 | HCF* | 32 | 8 | 19 | 4 | 1 | 0 | - | - | - |
| 29 | HCF 6 | 18 | 2 | 11 | 5 | 0 | 0 | 8 | 10 | 0 |
| 30 | HCF* | 24 | 5 | 10 | 8 | 1 | 0 | - | - | - |
| 31 | HCF 7 | 19 | 4 | 12 | 2 | 1 | 0 | 10 | 9 | 0 |
| 32 | HCF* | 21 | 5 | 13 | 3 | 0 | 0 | - | - | - |
| 33 | NHCF | 77 | 22 | 35 | 18 | 2 | 0 | - | - | - |
| 34 | HCF 8 | 18 | 5 | 6 | 7 | 0 | 0 | 9 | 9 | 0 |
| 35 | HCF 9 | 19 | 2 | 14 | 2 | 1 | 0 | 8 | 9 | 2 |
| 36 | HCF 21 | 32 | 10 | 14 | 6 | 2 | 0 | 18 | 14 | 0 |
| 37 | HCF* | 15 | 4 | 8 | 3 | 0 | 0 | - | - | - |
| 38 | HCF* | 23 | 10 | 11 | 2 | 0 | 0 | - | - | - |
| 39 | HCF* | 15 | 2 | 10 | 3 | 0 | 0 | - | - | - |
| 40 | HCF* | 12 | 6 | 2 | 3 | 1 | 0 | - | - | - |
| 41 | NHCF | 44 | 10 | 23 | 7 | 4 | 0 | - | - | - |
| 42 | HCF* | 8 | 4 | 3 | 1 | 0 | 0 | - | - | - |
| 43 | HCF* | 8 | 4 | 3 | 1 | 0 | 0 | - | - | - |
| 44 | HCF 10 | 9 | 4 | 3 | 2 | 0 | 0 | 5 | 4 | 0 |
| 45 | HCF 11 | 8 | 1 | 4 | 3 | 0 | 0 | 1 | 7 | 0 |
| 46 | HCF* | 2 | 0 | 2 | 0 | 0 | 0 | - | - | - |
| 47 | HCF 22 | 5 | 2 | 2 | 0 | 1 | 0 | 4 | 1 | 0 |
| All PCFs/HCFs |  | 2374 | 1463 | 716 | 174 | 20 | 1 | 1276 | 209 | 11 |

**Table S3.**
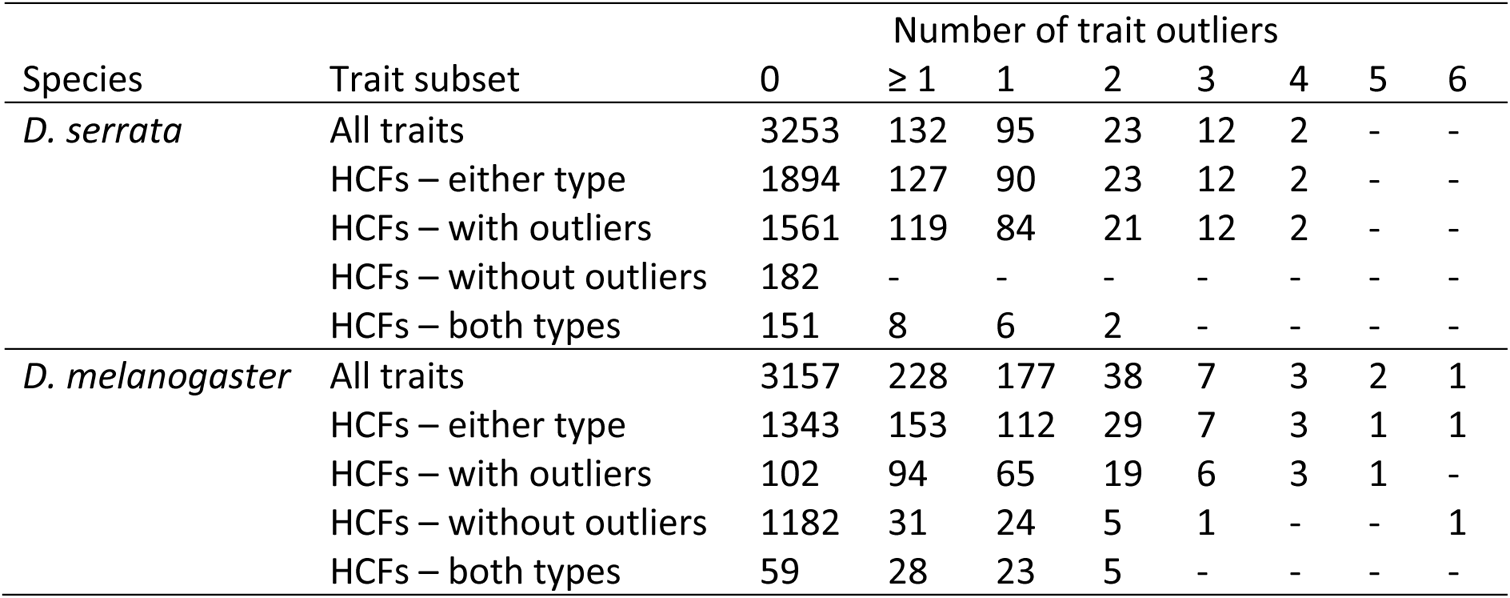
Count of expression traits with outlying lines for the two *Drosophila* gene expression datasets. Overall, 132 (3.9%) gene expression traits in *D. serrata* and 228 (6.7%) in *D. melanogaster* had at least one extreme value (≥ 1). Most of these traits had only a single outlier line but ranged up to four outlier lines (for two traits) in *D. serrata* and six outlier lines for one trait in *D. melanogaster*. We then subset the total 3,385 traits into those associated with HCFs (either type), then the specific category of HCFs, and finally traits that were shared between both types of HCF. For example, out of the 132 traits with outliers in *D. serrata*, 127 are associated with at least one HCF; 119 of these are associated only with one or more HCFs with outliers, none are associated only with an HCF without outliers, and eight are associated with at least one of each type of HCF.

| Species | Trait subset | Number of trait outliers |  |  |  |  |  |  |  |
| --- | --- | --- | --- | --- | --- | --- | --- | --- | --- |
|  |  | 0 | ≥ 1 | 1 | 2 | 3 | 4 | 5 | 6 |
| *D. serrata* | All traits | 3253 | 132 | 95 | 23 | 12 | 2 | - | - |
|  | HCFs – either type | 1894 | 127 | 90 | 23 | 12 | 2 | - | - |
|  | HCFs – with outliers | 1561 | 119 | 84 | 21 | 12 | 2 | - | - |
|  | HCFs – without outliers | 182 | - | - | - | - | - | - | - |
|  | HCFs – both types | 151 | 8 | 6 | 2 | - | - | - | - |
| *D. melanogaster* | All traits | 3157 | 228 | 177 | 38 | 7 | 3 | 2 | 1 |
|  | HCFs – either type | 1343 | 153 | 112 | 29 | 7 | 3 | 1 | 1 |
|  | HCFs – with outliers | 102 | 94 | 65 | 19 | 6 | 3 | 1 | - |
|  | HCFs – without outliers | 1182 | 31 | 24 | 5 | 1 | - | - | 1 |
|  | HCFs – both types | 59 | 28 | 23 | 5 | - | - | - | - |

**Table S4.** Distribution across chromosomes of ESTs associated with HCFs in the D. serrata lines. We mapped the 3385 analyzed ESTs to chromosomes of the reference *D. serrata* genome using BLASTn, resulting in “null” frequencies on X (446 ESTs), 2L (545), 2R (683), 3L (688), 3R (817) and “other” (206 ESTs that didn’t align to any of the five largest chromosomes). We then conducted chi-square tests of the chromosome frequencies of the subsets of traits associated with each individual HCF. HCFs with observed frequencies differing significantly from the null (s<.01) are shown in bold.

| | Chr X | | Chr 2L | | Chr 2R | | Chr 3L | | Chr 3R | | Other | | $\chi^2$ | p | s |
| --- | --- | --- | --- | --- | --- | --- | --- | --- | --- | --- | --- | --- | --- | --- | --- |
|  | Obs | Exp | Obs | Exp | Obs | Exp | Obs | Exp | Obs | Exp | Obs | Exp |  |  |  |
| 1 | 51 | 63.64 | 71 | 77.77 | 117 | 97.46 | 95 | 98.17 | 126 | 116.58 | 23 | 29.39 | 9.3 | 0.099 | 0.037 |
| 2 | 39 | 30.83 | 30 | 37.68 | 51 | 47.21 | 35 | 47.56 | 73 | 56.48 | 6 | 14.24 | 17 | 0.005 | 0.001 |
| 3 | 23 | 25.43 | 25 | 31.07 | 44 | 38.94 | 41 | 39.23 | 54 | 46.58 | 6 | 11.75 | 6.1 | 0.292 | 0.093 |
| 4 | 10 | 17.92 | 23 | 21.9 | 44 | 27.44 | 33 | 27.64 | 23 | 32.82 | 3 | 8.28 | 20.9 | 0.001 | <0.001 |
| 5 | 9 | 13.7 | 15 | 16.74 | 14 | 20.98 | 30 | 21.14 | 33 | 25.1 | 3 | 6.33 | 12.1 | 0.034 | 0.012 |
| 6 | 10 | 20.69 | 30 | 25.28 | 33 | 31.68 | 35 | 31.91 | 40 | 37.89 | 9 | 9.55 | 6.9 | 0.228 | 0.069 |
| 7 | 13 | 17.92 | 29 | 21.9 | 20 | 27.44 | 36 | 27.64 | 31 | 32.82 | 7 | 8.28 | 8.5 | 0.131 | 0.046 |
| 8 | 10 | 17 | 30 | 20.77 | 23 | 26.03 | 32 | 26.22 | 28 | 31.14 | 6 | 7.85 | 9.4 | 0.095 | 0.032 |
| 9 | 8 | 14.89 | 22 | 18.19 | 24 | 22.8 | 26 | 22.97 | 26 | 27.27 | 7 | 6.88 | 4.5 | 0.479 | 0.127 |
| 10 | 5 | 15.81 | 19 | 19.32 | 23 | 24.21 | 33 | 24.39 | 34 | 28.96 | 6 | 7.3 | 11.6 | 0.041 | 0.016 |
| 11 | 12 | 14.1 | 16 | 17.23 | 25 | 21.59 | 24 | 21.75 | 26 | 25.83 | 4 | 6.51 | 2.1 | 0.829 | 0.217 |
| 12 | 15 | 13.18 | 13 | 16.1 | 23 | 20.18 | 24 | 20.32 | 22 | 24.14 | 3 | 6.09 | 3.7 | 0.599 | 0.157 |
| 13 | 5 | 10.94 | 11 | 13.36 | 25 | 16.75 | 20 | 16.87 | 19 | 20.03 | 3 | 5.05 | 9.2 | 0.102 | 0.041 |
| 14 | 14 | 12.39 | 15 | 15.13 | 22 | 18.97 | 21 | 19.11 | 18 | 22.69 | 4 | 5.72 | 2.4 | 0.796 | 0.197 |
| 15 | 6 | 11.2 | 10 | 13.69 | 23 | 17.15 | 23 | 17.28 | 19 | 20.52 | 4 | 5.17 | 7.7 | 0.175 | 0.061 |
| 16 | 5 | 10.94 | 17 | 13.36 | 23 | 16.75 | 13 | 16.87 | 22 | 20.03 | 3 | 5.05 | 8.5 | 0.133 | 0.051 |
| 17 | 5 | 6.98 | 8 | 8.53 | 3 | 10.69 | 13 | 10.77 | 20 | 12.79 | 4 | 3.23 | 10.8 | 0.055 | 0.022 |
| 18 | 3 | 10.28 | 15 | 12.56 | 22 | 15.74 | 17 | 15.85 | 20 | 18.83 | 1 | 4.75 | 11.2 | 0.047 | 0.019 |
| 19 | 9 | 10.41 | 9 | 12.72 | 9 | 15.94 | 21 | 16.06 | 28 | 19.07 | 3 | 4.81 | 10.7 | 0.058 | 0.027 |
| 20 | 4 | 9.35 | 13 | 11.43 | 10 | 14.33 | 17 | 14.43 | 26 | 17.14 | 1 | 4.32 | 12.2 | 0.032 | 0.009 |
| 21 | 5 | 5.8 | 6 | 7.08 | 4 | 8.88 | 13 | 8.94 | 13 | 10.62 | 3 | 2.68 | 5.4 | 0.373 | 0.114 |
| 22 | 0 | 3.69 | 3 | 4.51 | 2 | 5.65 | 17 | 5.69 | 2 | 6.76 | 4 | 1.7 | 35.5 | <0.001 | <0.001 |
| 23 | 3 | 3.95 | 5 | 4.83 | 13 | 6.05 | 6 | 6.1 | 3 | 7.24 | 0 | 1.83 | 12.5 | 0.028 | 0.006 |
| 24 | 2 | 2.9 | 6 | 3.54 | 3 | 4.44 | 5 | 4.47 | 5 | 5.31 | 1 | 1.34 | 2.6 | 0.759 | 0.177 |
| 25 | 0 | 2.77 | 2 | 3.38 | 5 | 4.24 | 4 | 4.27 | 9 | 5.07 | 1 | 1.28 | 6.6 | 0.253 | 0.084 |
| 26 | 0 | 6.06 | 3 | 7.41 | 4 | 9.28 | 1 | 9.35 | 3 | 11.1 | 35 | 2.8 | 395.4 | <0.001 | <0.001 |
| 27 | 12 | 11.33 | 18 | 13.85 | 19 | 17.35 | 12 | 17.48 | 22 | 20.76 | 3 | 5.23 | 4.2 | 0.523 | 0.141 |
| 28 | 21 | 18.71 | 26 | 22.86 | 24 | 28.65 | 31 | 28.86 | 37 | 34.27 | 3 | 8.64 | 5.5 | 0.355 | 0.103 |
| 29 | 4 | 6.98 | 11 | 8.53 | 10 | 10.69 | 18 | 10.77 | 10 | 12.79 | 0 | 3.23 | 10.7 | 0.057 | 0.025 |
| 30 | 1 | 0.79 | 1 | 0.97 | 1 | 1.21 | 1 | 1.22 | 1 | 1.45 | 1 | 0.37 | 1.4 | 0.927 | 0.239 |
| 31 | 0 | 1.32 | 0 | 1.61 | 0 | 2.02 | 0 | 2.03 | 0 | 2.41 | 10 | 0.61 | 154.3 | <0.001 | <0.001 |
| 32 | 0 | 0.53 | 1 | 0.64 | 0 | 0.81 | 0 | 0.81 | 3 | 0.97 | 0 | 0.24 | 6.9 | 0.23 | 0.077 |

**Table S5.**
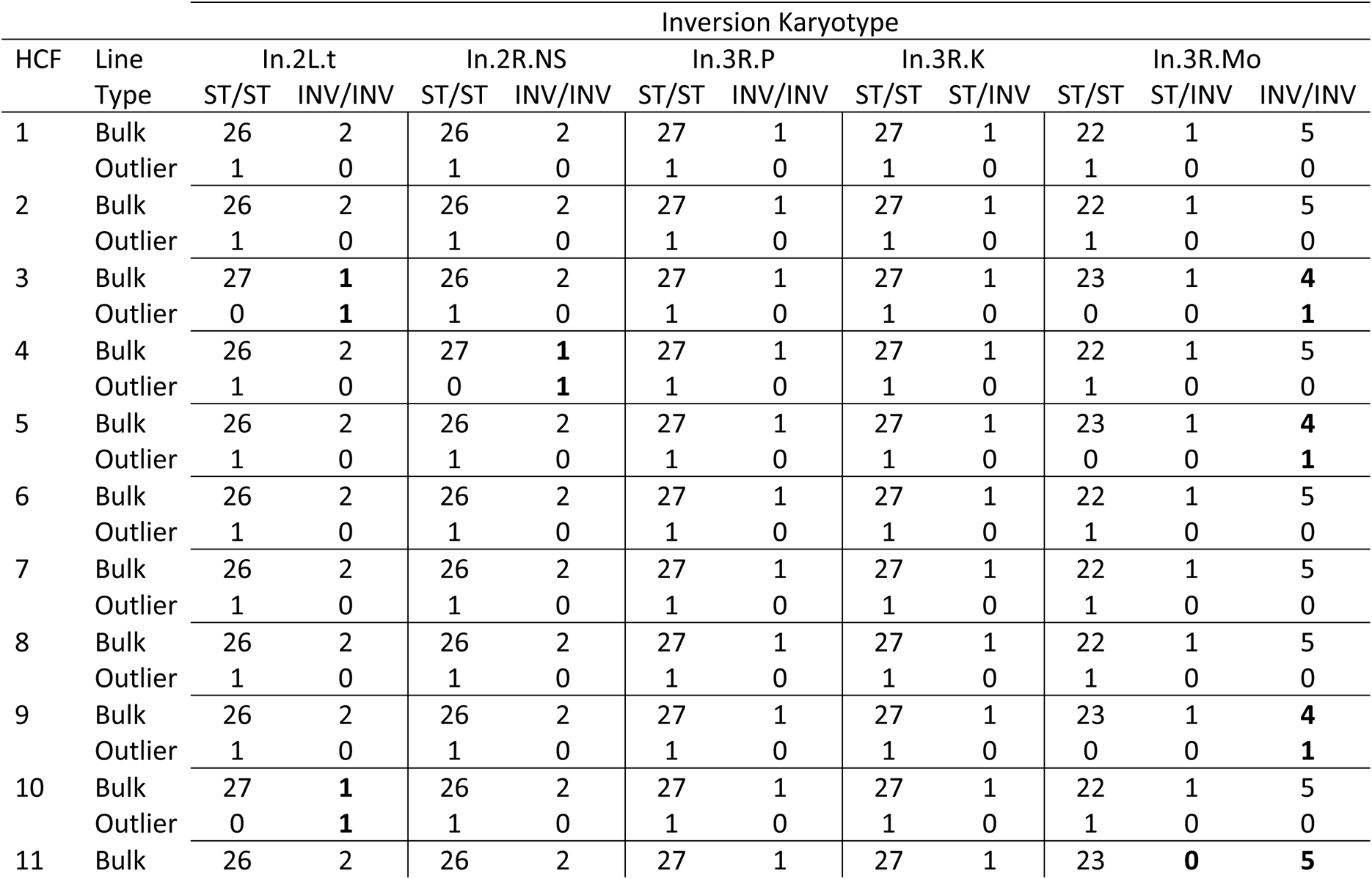
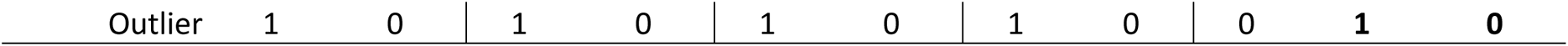
Karyotypes for five inversions and HCF score outlier status for 29 of the 30 *D. melanogaster* lines. Five known inversions were segregating as one or two copies across 11 of the 29 lines that were included in the BSFG analysis and had inversion karyotypes. All outlier lines for HCFs 1, 2 and 6-8 had the standard karyotype (ST/ST) for each of the five inversions. The outlier line for each of HCFs 3, 4, 5 and 9-11 carried at least one copy (kayrotypes ST/INV or INV/INV) of one (HCFs 4,5,9-11) or two (HCF 3) types of inversion. In each of these cases, at least one line falling within the bulk of the distribution for that HCF also carried at least one copy of the inversion (shown in bold).

**Table S6.**
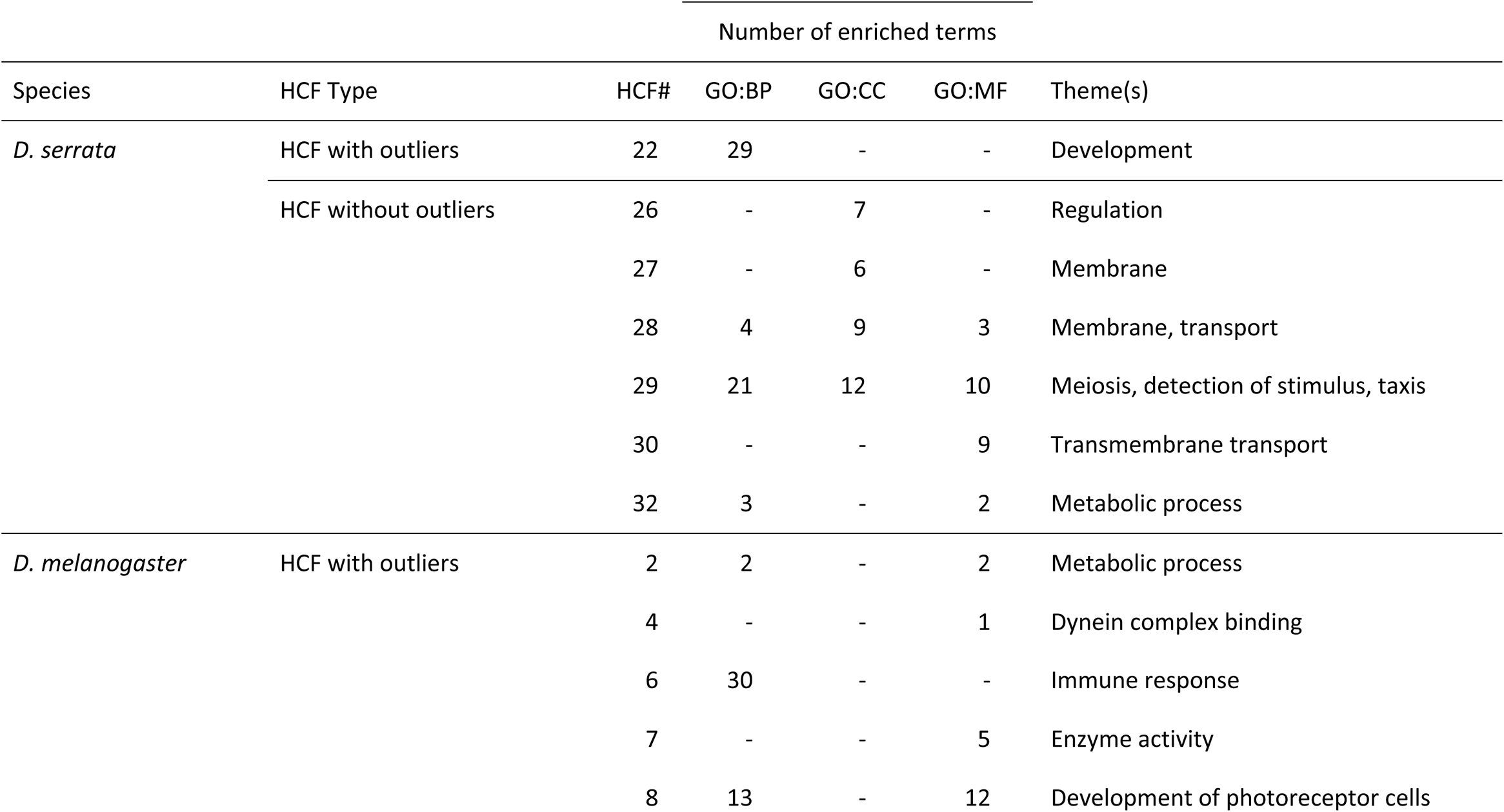
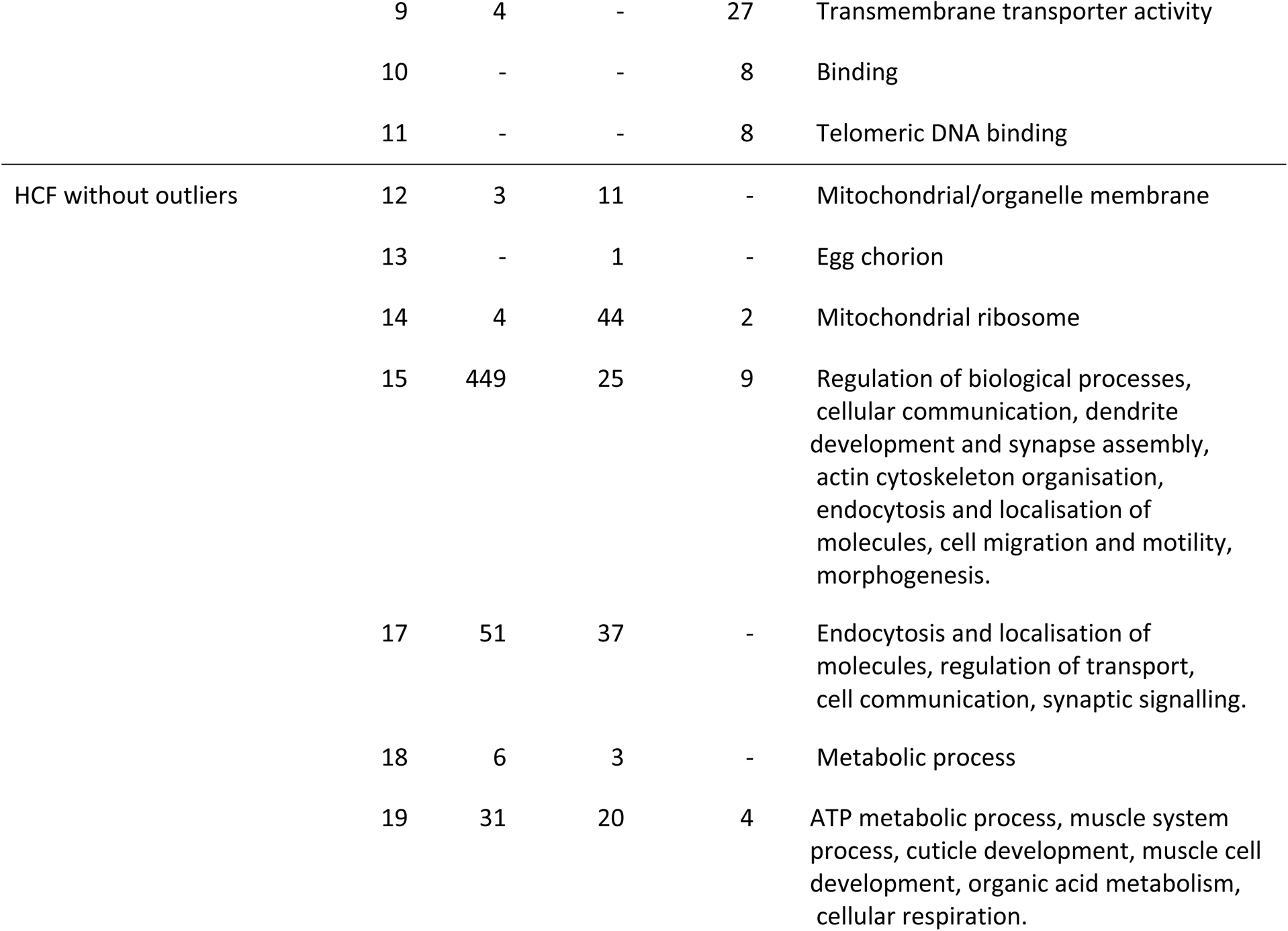
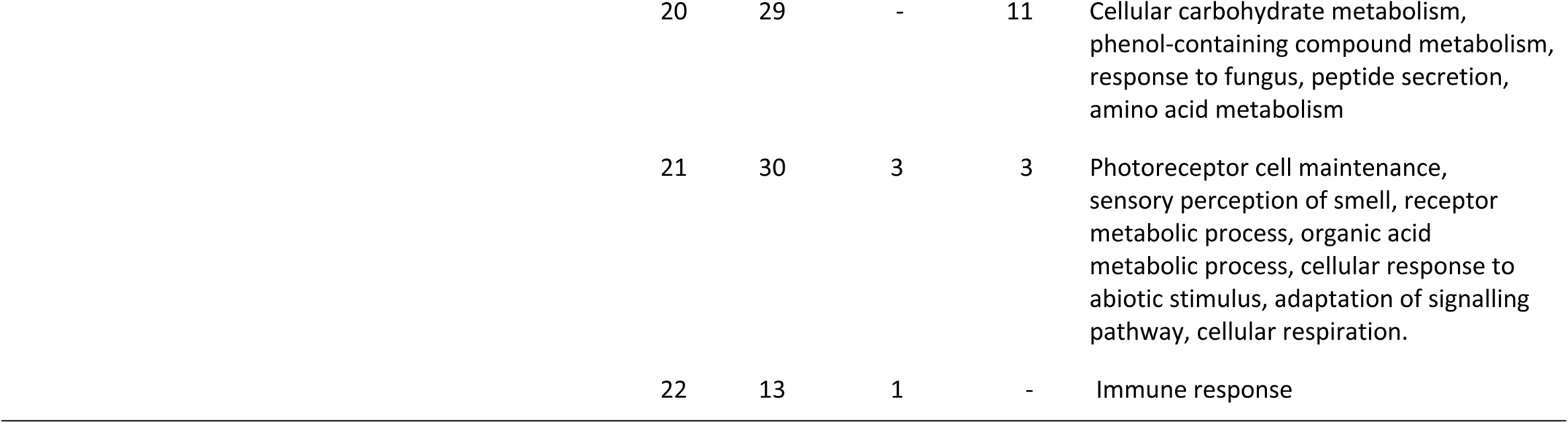
Summary of functional enrichment analyses conducted using g:Profiler and semantic similarity analysis conducted with GOSemSim. Only HCFs with significant enrichment for at least one term in any of the three Gene Ontology (GO) categories are shown. The full lists of significantly enriched terms can be downloaded at https://doi.org/10.48610/90441fc.

**Figure S1.**
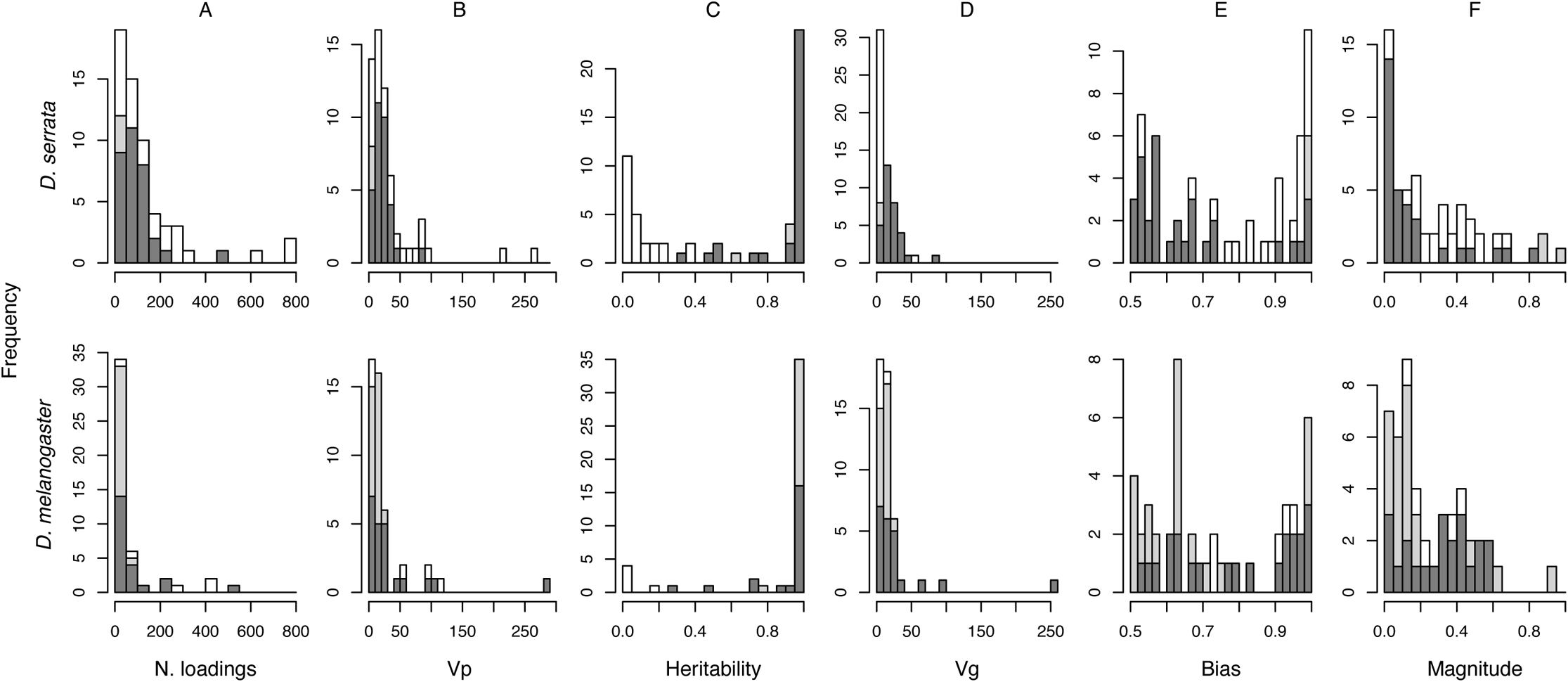
Characteristics of phenotypic common factors (PCFs) estimated in Bayesian Sparse Factor Genetic (BSFG) analyses of gene expression data from two *Drosophila* datasets. We used several metrics (columns A to F) to characterize and compare the 59 *D. serrata* (top row) and 47 *D. melanogaster* (bottom row) estimated PCFs, by PCF type. Black (grey) bars indicate the frequencies of heritable PCFs that were (were not) investigated further (see methods, Fig S2), while white bars indicate non-heritable PCF frequencies. The maximum (771 vs 549) and median (83 vs 33) number of significant trait loadings were substantially larger for *D. serrata* PCFs than *D. melanogaster* (column A). PCFs accounted for 59% of the phenotypic variance in *D. serrata* compared to 39% in *D. melanogaster*, reflecting both the larger number of PCFs in *D. serrata* and the presence of two PCFs with relatively large phenotypic variance compared to only one in *D. melanogaster* (column B). PCF heritabilities were lower on average in *D. serrata* than *D. melanogaster* (column C) with medians of 0.73 and 0.99 respectively. This translated to PCFs accounting for less genetic variance (column D) in *D. serrata* than *D. melanogaster*, with medians of 7.42 and 11.53 and totals of 779.82 and 957.28 across PCFs in *D. serrata* and *D. melanogaster*, respectively. There was no clear difference between the species, or between heritable and non-heritable PCFs in the proportion of significant trait loadings acting in the same direction as the median significant trait loading for a given PCF (i.e., bias: column E) or in the mean absolute value of significant trait loadings (i.e., magnitude: column F).

**Figure S2.**
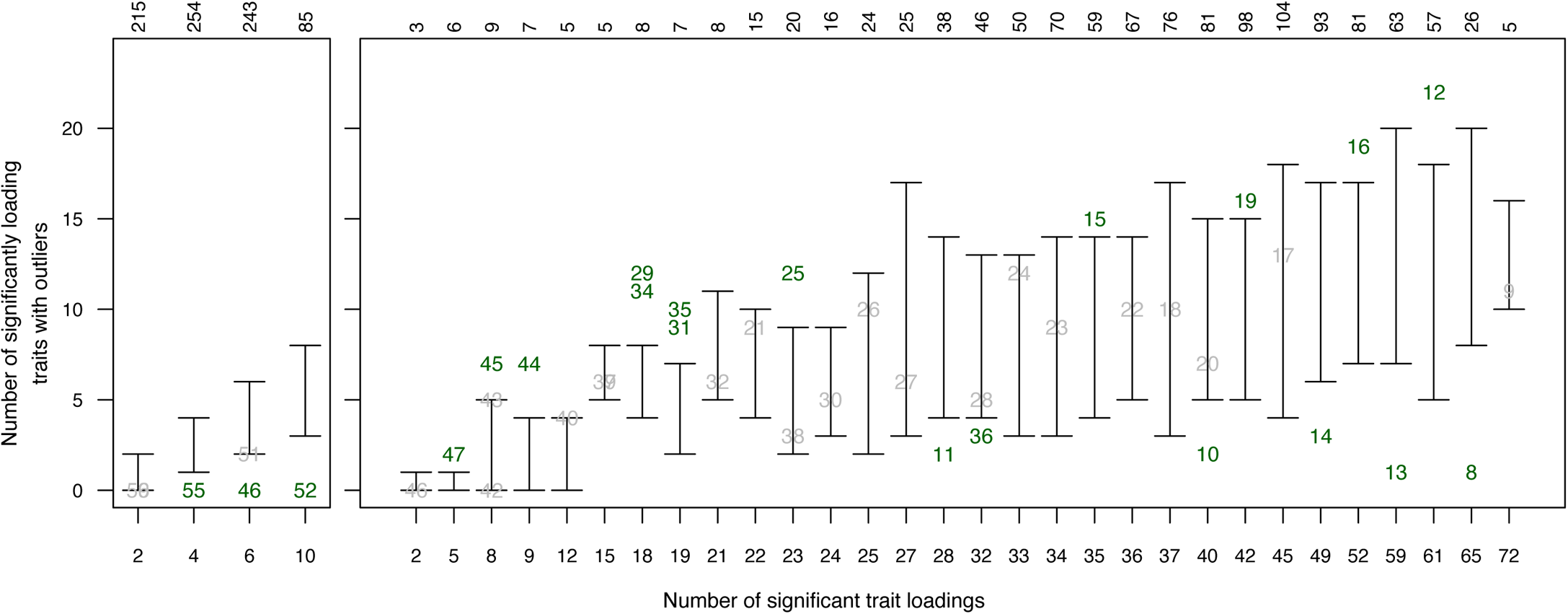
Influence of individual trait outliers on HCF estimation in randomized datasets, and comparison to observed data for *D. serrata* (A) and *D. melanogaster* (B). For a given observed HCF (Tables S1–S4), we first identified the subset of randomized data HCFs (number of randomized data HCF in that subset is shown on top x-axis) with the same number of significant trait loadings (bottom x-axis). We then compared the number of significantly loading traits with outlying lines (y-axis) between the observed HCF and the matched subset of randomized data HCFs. We excluded from subsequent investigation any observed HCFs (labelled, in grey, by PCF number reported in Table S1 or S2) that fell within the range of significantly loading traits with outliers that was typically observed in the randomized data HCFs. Where the number of outlying-line traits for the observed HCF fell outside the 95% range of the randomized data HCF subset, we retained the observed HCF for analysis (labelled by PCF number in green). We thereby excluded three *D. serrata* HCFs (note, PCFs 56 and 59, each with two significant trait loadings, are superimposed) and 20 *D. melanogaster* HCFs (note, PCFs 37 and 39, with 15 significant trait loadings, are superimposed) from further investigation.

**Figure S3.**
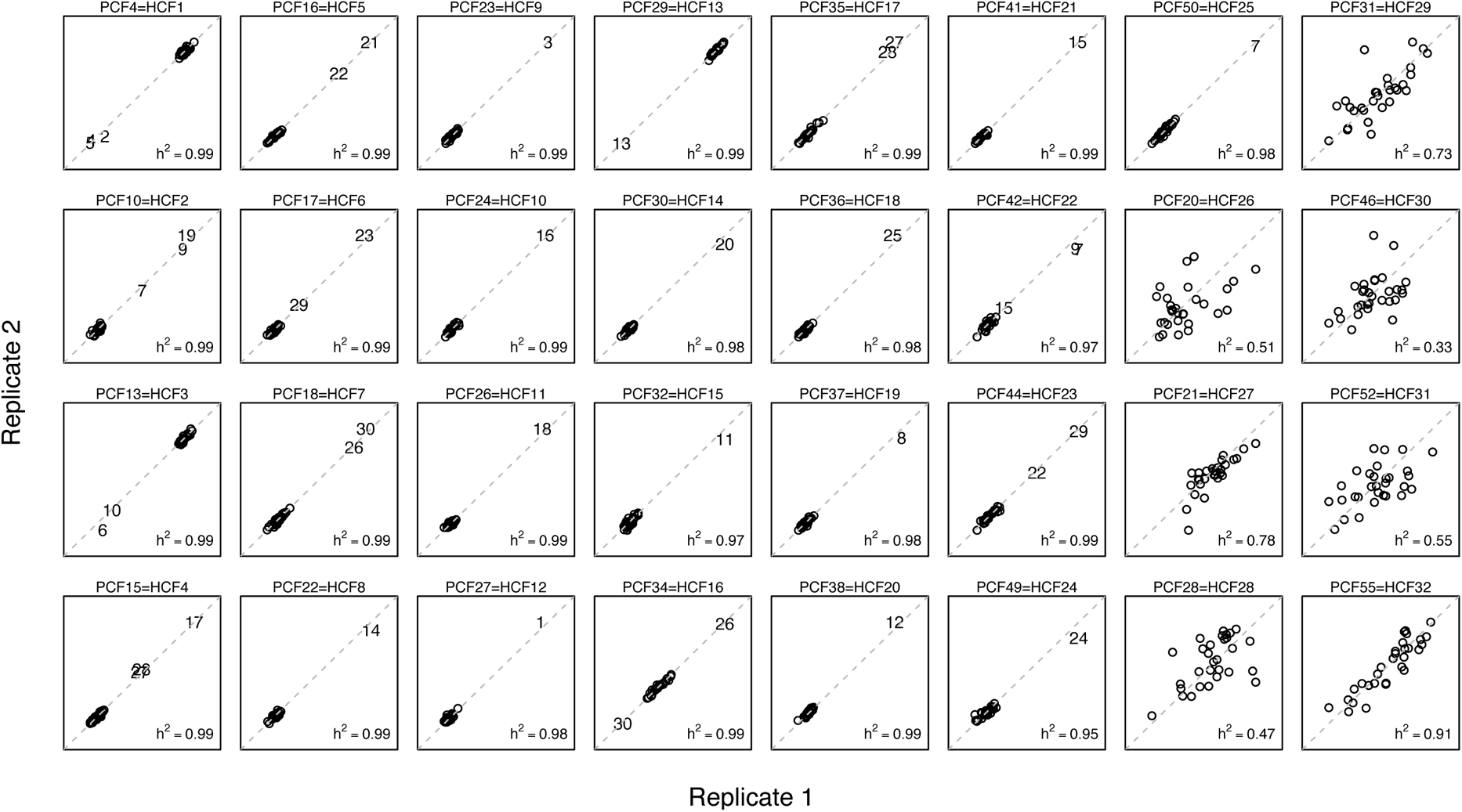
PCF scores for replicate pairs of the 30 *D. serrata* lines for the 32 heritable phenotypic common factors. Replicate-level PCF scores are presented, with the two replicates per line designated replicate 1 and replicate 2 arbitrarily. Each graph is labeled above with the PCF and HCF numbers, as shown in Table S1, and are ordered first by HCF type (HCF1 to HCF25 have outlier line(s), while HCF26 to HCF32 do not) then by decreasing contribution to phenotypic variance. Numbered values indicate individual inbred lines that met the criteria for being outliers (see Methods) on that HCF; notably, the same individual line was not an outlier across many HCF (i.e., the same line number is not consistently repeated across panels). The dashed line indicates a 1:1 relationship between line replicate PCF scores, where tighter clustering of PCF scores along this line is associated with higher heritability (heritability shown in the bottom left of each panel), and low replicate-specific (non-genetic) influences.

**Figure S4.**
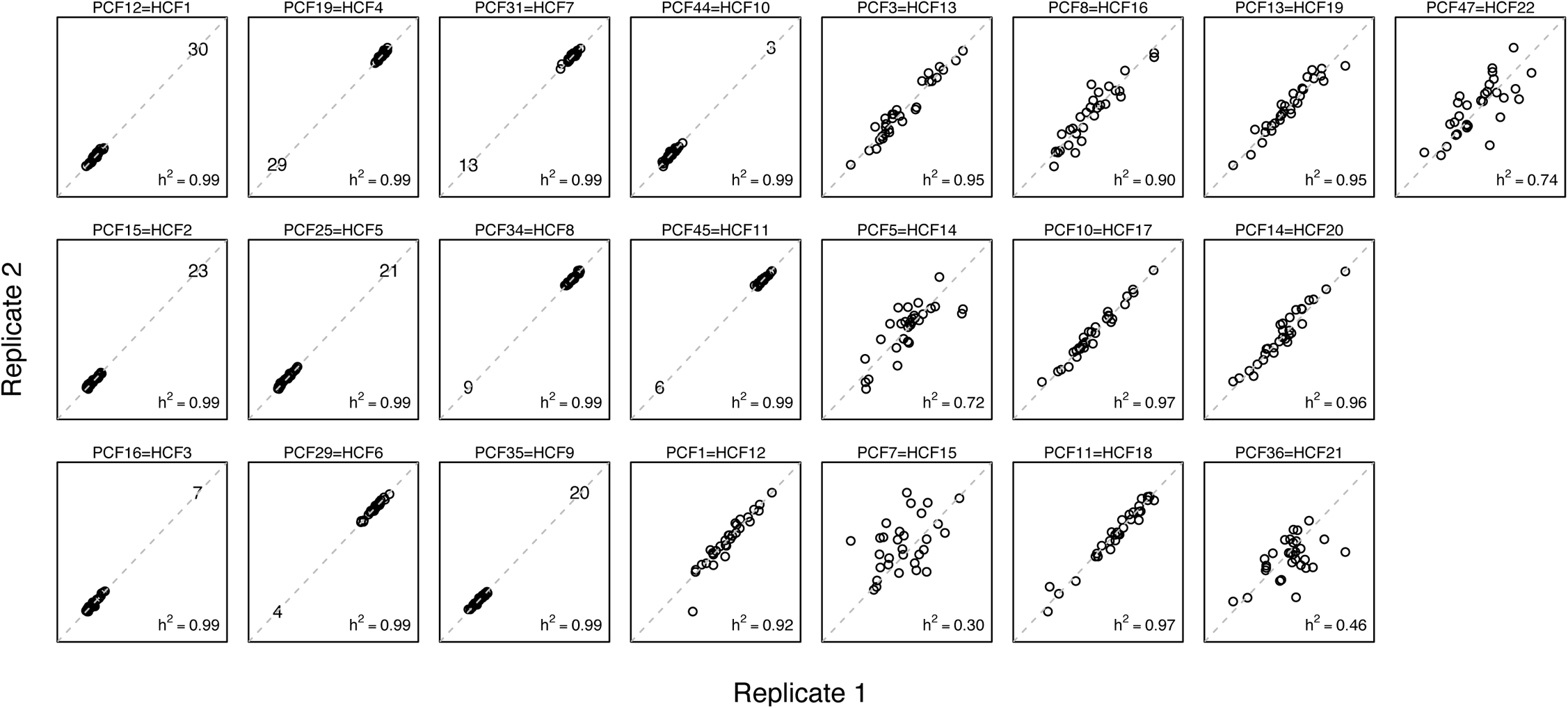
PCF scores for replicate pairs of the 30 D. melanogaster lines for the 22 heritable phenotypic common factors. Replicate-level PCF scores are presented, with the two replicates per line designated replicate 1 and replicate 2 arbitrarily. Each graph is labeled above with the PCF and HCF numbers, as shown in Table S1, and are ordered first by HCF type (HCF1 to HCF11 have outlier line(s), while HCF12 to HCF22 do not) then by decreasing contribution to phenotypic variance. Numbered values indicate individual inbred lines that met the criteria for being outliers (see Methods) on that HCF; notably, the same individual line was not an outlier across many HCF (i.e., the same line number is not consistently repeated across panels). The dashed line indicates a 1:1 relationship between line replicate PCF scores, where tighter clustering of PCF scores along this line is associated with higher heritability (heritability shown in the bottom left of each panel), and low replicate-specific (non-genetic) influences.

**Figure S5.**
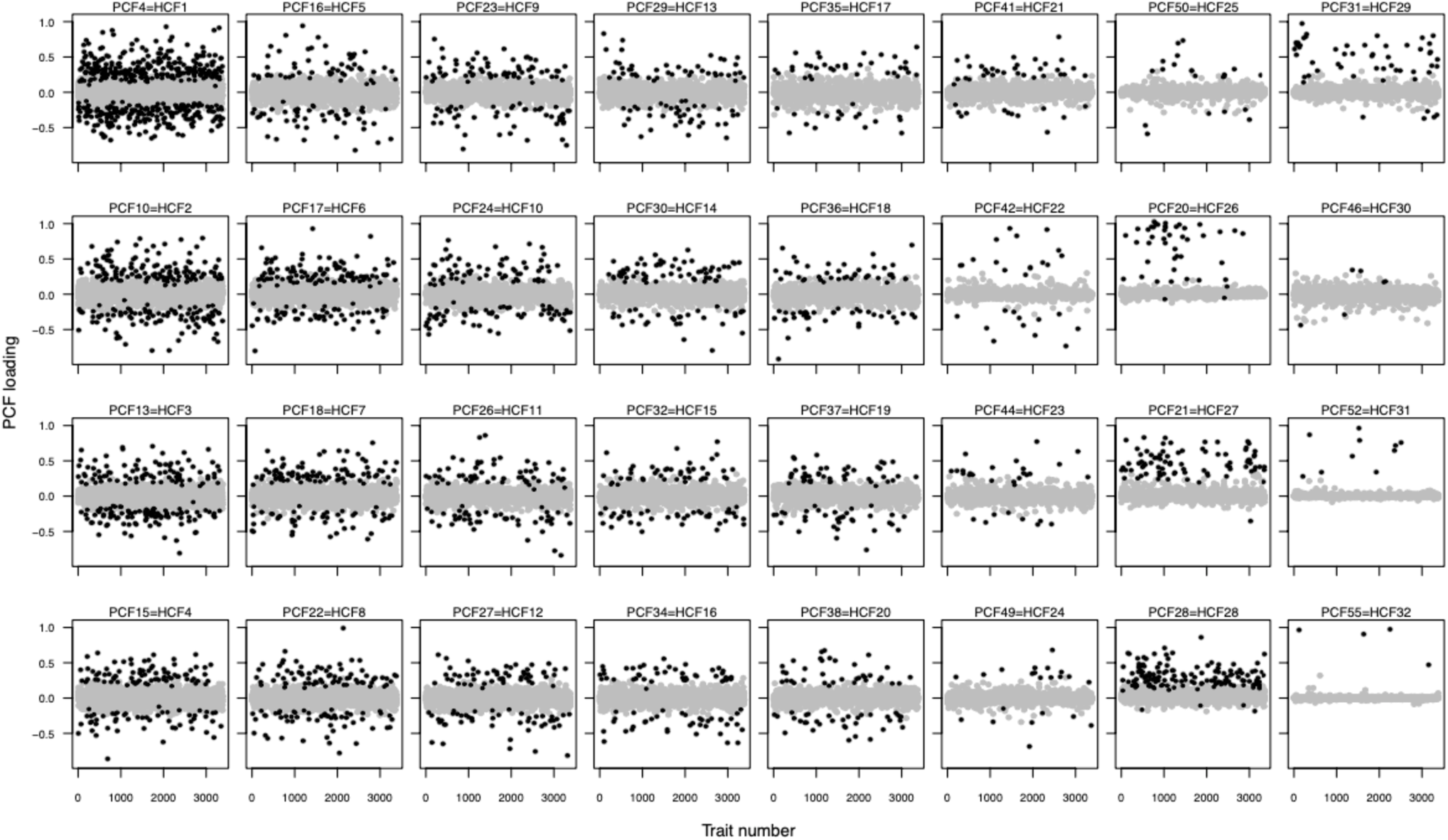
Trait loadings of the 32 *D. serrata* HCFs. Significant (nonsignificant) trait loadings are represented by black (grey) circles. Traits are plotted in order of EST number, where traits were arbitrarily assigned a numerical identifier prior to analyses. HCFs exhibiting the greatest degree of directional bias in trait loadings (HCFs 26-32) are also those without outlier lines (Fig. S3).

**Figure S6.**
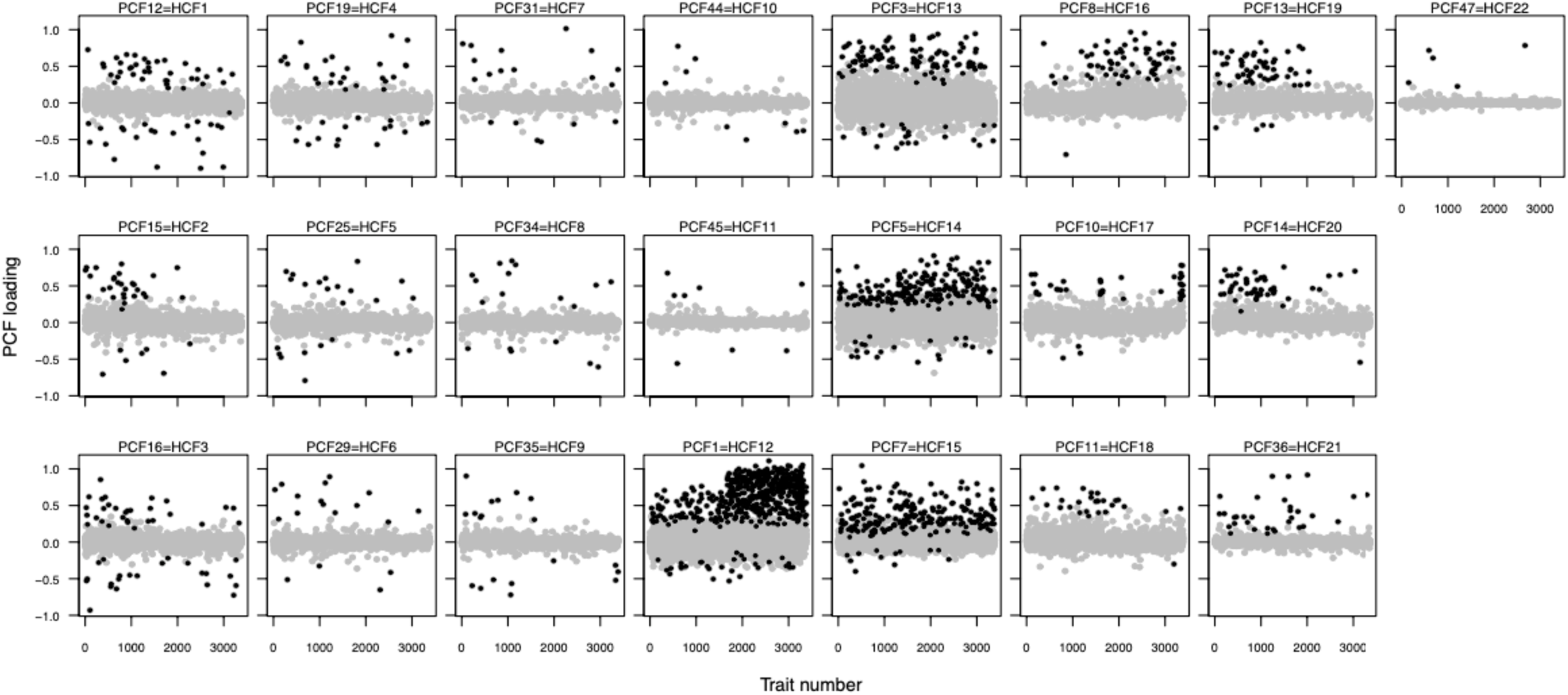
Trait loadings of the 22 *D. melanogaster* HCFs. Significant (nonsignificant) trait loadings are depicted as black (grey) circles. Traits are plotted as ordered in the full dataset, which was arranged by module membership (D. Runcie, pers. comm). Notably, this ordering of traits reveals apparent relationships between the HCFs identified here, and co-associating sets of expression traits previously identified by Ayroles et al. (2009). Specifically, HCFs 12 and 16 appear to relate strongly to one of the two large modules detected in the original gene expression analysis (i.e., higher numbered traits contribute), while HCFs 2, 19 and 20 appear to be strongly associated with the other (i.e., lower numbered traits contribute). As we observed for *D. serrata*, HCFs with the largest degree of directional bias (HCFs 12-22) are also those without outlier lines (Fig. S4).

